# Spaced training enhances memory and prefrontal ensemble stability in mice

**DOI:** 10.1101/2020.12.17.417451

**Authors:** Annet Glas, Mark Hübener, Tobias Bonhoeffer, Pieter M. Goltstein

## Abstract

Memory is substantially improved when learning is distributed over time, an effect called “spacing effect”. So far it has not been studied how spaced learning affects neuronal ensembles presumably underlying memory. In the present study, we investigate whether trial spacing increases the stability or size of neuronal ensembles. Mice were trained in the “everyday memory” task, an appetitive, naturalistic, delayed matching-to-place task. Spacing trials by 60 minutes produced more robust memories than training with shorter or longer intervals. c-Fos labeling and chemogenetic inactivation established the necessity of the dorsomedial prefrontal cortex (dmPFC) for successful memory storage. *In vivo* calcium imaging of excitatory dmPFC neurons revealed that longer trial spacing increased the similarity of the population activity pattern on subsequent encoding trials and upon retrieval. Conversely, trial spacing did not affect the size of the total neuronal ensemble or the size of subpopulations dedicated to specific task-related behaviors and events. Thus, spaced learning promotes reactivation of prefrontal neuronal ensembles processing episodic-like memories.

## Introduction

Extending the period between individual learning events can considerably strengthen a memory and increase its lifespan, a phenomenon called the “spacing effect” ^1^. This phenomenon has been described across a wide range of species, from mollusk to man ^2^. In mice, spaced training can strengthen associative ^3^ episodic-like ^4^, motor ^5^, and spatial ^6^ memories. The effectiveness of spacing learning is thought to be mediated by molecular and synaptic processes ^2^, which involve activation and expression of key signaling proteins and transcription factors ^2; 6; 7^, leading to increased synaptic plasticity ^5; 8^. It has not yet been studied, whether and how increasing the spacing of learning events affects neuronal ensembles representing an individual memory.

During a learning experience, a subset of neurons is activated as a result of their intrinsic excitability and external sensory drive ^9; 10; 11; 12^. The memory itself is thought to be encoded by synaptic connections that are newly formed or strengthened within this neuronal ensemble ^13; 14; 15^. Subsequently, memories can be consolidated by further functional and structural synaptic remodeling, enabling long-term retention ^16; 17^. For retrieval of a memory, neurons that are part of the ensemble need to be reactivated in a pattern similar to that during memory encoding ^11; 18; 19^.

The working hypothesis for the present work is that the molecular and synaptic mechanisms underlying the spacing effect ^2^ can influence two characteristics of neuronal ensembles, i.e. the size or reactivation pattern of the ensemble, during memory encoding, storage, and retrieval. The reasoning is that when learning occurs over multiple optimally spaced trials, molecular signaling initiated in the first trial can extend the temporal window of enhanced neuronal excitability ^10; 20^ and thereby increase the likelihood of the same ensemble being reactivated in subsequent trials. As such, spaced training would more effectively strengthen the ensemble’s internal synaptic connectivity ^21; 22^ and by local competitive circuit interactions result in a sparser, but more reliably activated assembly ^23; 24; 25^. Sparseness would safeguard the specificity of the represented memory ^26^, while stronger connectivity would render the memory more resilient to homeostatic mechanisms that can result in forgetting ^27^ and thereby increase the probability of retrieval. Conversely, as the group of excitable neurons drifts over time ^10^, consecutive learning experiences could activate different sets of neurons. Spacing learning experiences over extended periods could therefore allocate a memory to overlapping sets of neurons ^28^. Within this framework, the memory enhancing effect of spaced training could be mediated by representing a learning experience with a larger neuronal ensemble ^4; 11^.

To determine whether and how trial spacing changes the way neuronal ensembles represent learned experiences, we implemented the “everyday memory” task, a naturalistic delayed matching-to-place task ^7^. The instilled episodic-like memories are typically forgotten within 24 hours, but spaced training reliably prolongs the period over which the memories can be retrieved ^7^. Efficient execution of the everyday memory task relies on functions that have been attributed to the dorsal medial prefrontal cortex (dmPFC), including behavioral flexibility ^29^ and learning against a background of relevant prior knowledge ^30^. Moreover, rodent prefrontal cortex is, in concert with hippocampus, involved in the encoding and retrieval of episodic-like memories ^31; 32^, providing an attractive system for examining the relation between neuronal ensemble activity and memory strength.

Here we report that trial spacing improves memory and is accompanied with enhanced reactivation of the neuronal ensembles in dmPFC. Increasing trial spacing in the everyday memory task enhanced memory retrieval, yet impaired memory encoding. *In vivo* calcium imaging with a miniaturized microscope revealed that trial spacing results in more similar reactivation of the ensemble between encoding trials and upon memory retrieval. Conversely, trial spacing did not affect the size of the ensemble, suggesting that trial spacing primarily affects the synaptic strength within the neuronal ensemble but not its size.

## Results

### Studying episodic-like memory in an “everyday memory” task

We trained female mice (n = 20) in repeated sessions of an “everyday memory” task (see Method Summary, **Figure 1A, B, Video 1**; ^7^). Each training session consisted of three encoding trials (ETs; separated by an “encoding intertrial interval”) and three retrieval trials (RTs). During each encoding trial, mice entered the radial arm maze from a start box, explored the maze and retrieved a buried chocolate reward by digging in one of two available, odor-masked sandwells (i.e. the “rewarded” sandwell; **Figure S1A–C**). Upon completion of the final encoding trial, mice were kept in their home cage for an extended delay period (“retrieval delay”), after which the three retrieval trials (RTs) were conducted. During retrieval trials, mice had to revisit the previously rewarded sandwell. Simultaneously, mice had to refrain from digging at the previously non-rewarded sandwell, as well as four new non-rewarded sandwells (“non-cued” sandwells).

**Figure 1.**
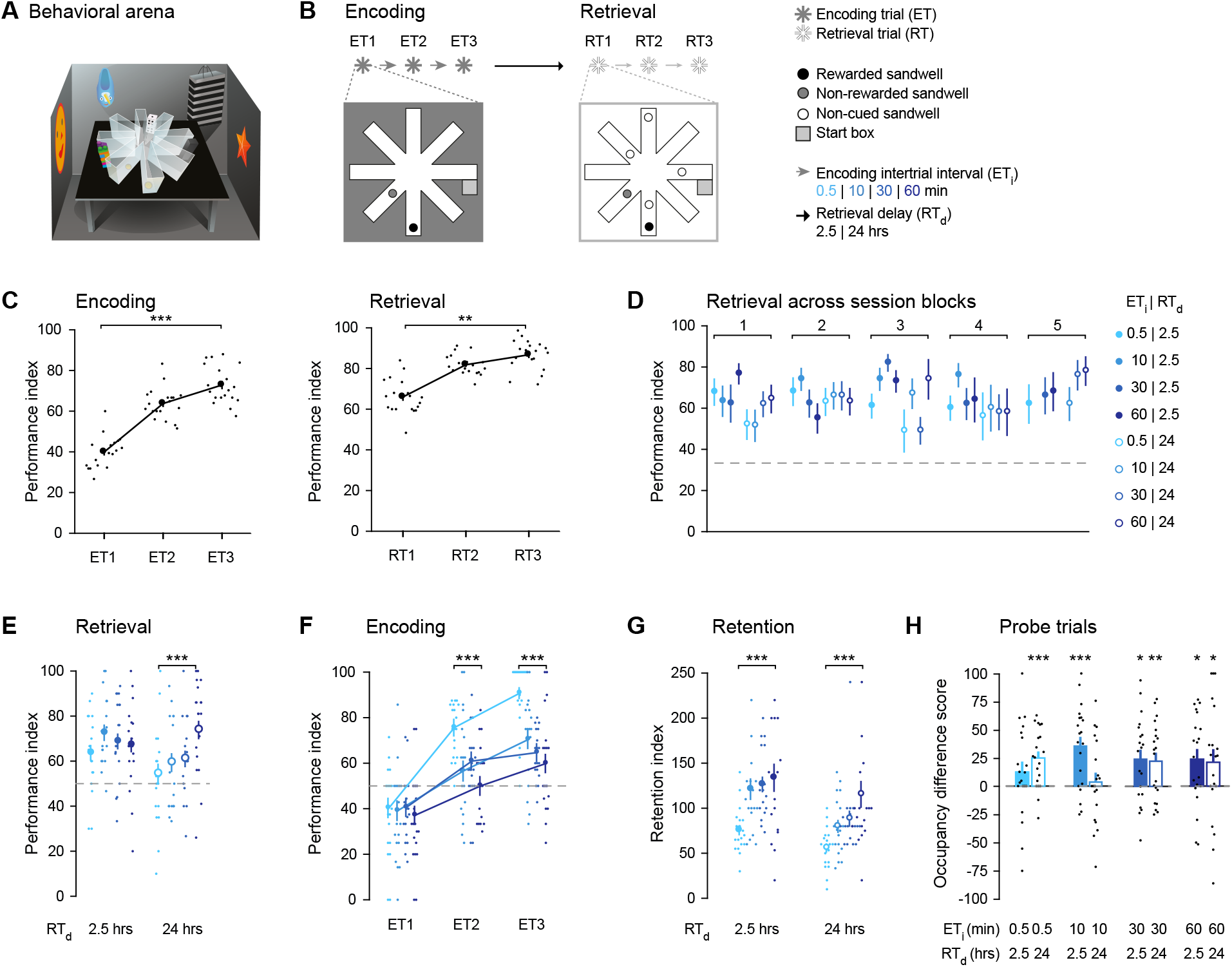
Trial spacing enhances memory retrieval, yet impairs memory encoding on the everyday memory task. **A** Schematic of behavioral training setup. **B** Schematic of session structure. Sandwell locations were altered on each session. **C** Performance improved across encoding (left) and retrieval trials (right; ET1 vs. RT3: X^2^_5_ = 75.6, p = 7.03·10^−15^, n = 19 mice; ET1 vs. ET3: T = −0.88, p = 2.38·10^−4^; RT1 vs. RT3: T = −0.62, p = 1.50·10^−3^, n = 19 mice). **D** Performance on RT1 remained stable over two months of behavioral training (first vs. last block of 8 sessions: U = 2.50·10^4^, p = 0.089). **E** Increasing the ET_i_ did not alter memory retrieval performance on RT1 after a 2.5-hrs RT_d_ (r_s_ = 0.06, p = 0.608), yet did after 24 hrs (r_s_ = 0.32, p = 5.67·10^−3^). **F** Increasing the ET_i_ impaired performance at ET2 (r_s_ = −0.44, p = 9.34·10^−5^, n = 19 mice) and ET3 (r_s_ = −0.53, p = 8.69·10^−7^, n = 19 mice). **G** Increasing the ET_i_ enhanced memory retention after a 2.5-hrs RT_d_ (r_s_ = 0.42, p = 2.70·10^−4^, n = 19 mice) and after a 24-hrs RT_d_ (r_s_ = 0.50, p = 7.63·10^−6^, n = 19 mice). **H** Memory was not observed after 2.5 hrs if training was conducted with an ETi of 0.5 min, but was after 24 hrs (ET_i_ 0.5 min, RT_d_ 2.5 hrs vs. chance: t_19_ = 1.35, p = 0.194; ET_i_ 0.5 min, RT_d_ 24 hrs vs. chance: t_19_ = 4.27, p = 4.12·10^−4^). Conversely, training with an ET_i_ of 10 min resulted in memory after 2.5 hrs, but not after 24 hrs (ET_i_ 10 min, RT_d_ 2.5 hrs vs. chance: t_19_ = 4.30, p = 3.89·10^−4^; ET_i_ 10 min, RT_d_ 24 hrs vs. chance: t_19_ = 0.43, p = 0.675). Memory was present and stable on probe trials conducted after training using a 30-min or 60-min ET_i_ (ET_i_ 30 min, RT_d_ 2.5 hrs vs. chance: t_19_ = 2.83, p = 0.011; ET_i_ 30 min, RT_d_ 24 hrs vs. chance: t_19_ = 2.94, p = 0.008; ET_i_ 60 min, RT_d_ 2.5 hrs vs. chance: t_19_ = 2.63, p = 0.017; ET_i_ 60 min, RT_d_ 2.5 hrs vs. chance: t_19_ = 2.48, p = 0.023). Filled dots indicate data from one mouse, circles and bars indicate mean (±SEM) across mice, and gray dashed lines indicates chance level. * p < 0.05, ** p < 0.01, *** p < 0.001.

After each session, we changed the spatial configuration of the sandwells and the position of the start box. Consequently, mice had to relearn and remember a different rewarded location in each subsequent session (**Figure S1D**). Performance in each trial was quantified as the number of incorrect sandwells the mouse dug in, relative to the total number of available sandwells (see Supplemental Information).

We first characterized the conditions under which mice were able to successfully complete the task. Memory was only reliably retrieved after training with multiple encoding trials in which the rewarded location was kept constant (**Figure S2**). In sessions, performance increased across subsequent encoding trials, as well as subsequent retrieval trials, verifying that mice can encode and retrieve memories in this task (**Figure 1C**). In addition, we studied the within- and between-session strategies that mice employ in this task. Altering the start box location between and after encoding trials confirmed that mice primarily used an allocentric (world-centered) rather than egocentric (body-centered) reference frame (**Figure S3A–D**; ^33^). Within a session, mice focused their search progressively closer to the rewarded arm (**Figure S3E**) and revisited non-rewarded arms less than expected from chance (**Figure S3F**). Between sessions, the previous session’s retrieval performance did not affect the next session’s retrieval performance (**Figure S3G**), suggesting that a successfully stored memory did not interfere with learning of a new memory. From these analyses, we conclude that mice employ both a “within-session win-stay” and “between-session switch” strategy to optimize their task performance.

### Increasing trial spacing enhances memory retrieval but impairs memory encoding

To examine the influence of trial spacing on encoding and retrieval of episodic-like memory, we tested the effect of four encoding intertrial intervals: 30 s (i.e. “massed” training, 119 sessions), 10 min (115 sessions), 30 min (133 sessions), and 60 min (132 sessions; **Figure 1B, Video 1**). To probe the effect of trial spacing on same- and next-day memory retrieval separately, we conducted retrieval trials after a retrieval delay of either 2.5 or 24 hrs. Performance was stable over months of training, allowing us to average a mouse’s performance across sessions of the same encoding intertrial interval and retrieval delay (**Figure 1D**).

Memory retrieval after 24 hrs was improved when encoding intertrial intervals were longer, yet no effect was observed after 2.5 hrs (**Figure 1E**). This difference was not unexpected as trial spacing primarily affects less recent memories ^2^. In addition, we observed that performance in the second and third encoding trial was reduced when encoding intertrial intervals were extended (**Figure 1F**). As a control, we compensated for impaired encoding by normalizing the performance in the first retrieval trial to the encoding performance in the final, third encoding trial (“retention”). Retention thereby addressed how much of the successfully encoded information persisted and subsequently could be retrieved. Memory retention positively correlated with encoding intertrial interval after both a 2.5 hrs and 24 hrs retrieval delay (**Figure 1G**).

In a subset of sessions, we quantified the absolute strength of the memory by conducting a probe trial, which replaced the first retrieval trial. In these probe trials, the previously rewarded sandwell did not contain reward for the first minute of exploration (**Video 2, 3**). Memory in probe trials was quantified as the relative dig time at the rewarded sandwell, normalized to the total dig time at the rewarded and non-rewarded sandwell (termed the “occupancy difference score”). In sessions conducted with spaced encoding intertrial intervals, we observed an inverted U-shaped effect of trial spacing on next-day memory. Specifically, mice that were trained using a 10 or 180 min encoding intertrial interval did not remember the rewarded location after 24 hrs (**Figure 1H, S4**), while memories persisted after training with encoding intertrial intervals of 30 min or 60 min (**Figure 1H**). Unexpectedly, massed training did not result in same-day memory, but memory was observed after 24 hrs and was even still present after 48 hrs (see Discussion, **Figure 1H, S4**).

Differences in trial spacing could affect a number of memory-related behavioral variables besides error-based performance: the latency to find the rewarded sandwell, distance traveled, running speed, relative dig time, and number of arm visits (**Figure S5**). In consecutive encoding trials, we observed a quantitative reduction in the variables that are indicative of exploration, i.e. latency, distance traveled, running speed, and number of arms visited. Conversely, we observed an increase in the relative dig time, a measure of exploitation of memory of the rewarded location. These results suggest that mice explored less and increasingly used their recollection of the rewarded sandwell location in subsequent encoding trials. However, none of these behavioral variables were significantly influenced by trial spacing. We conclude that increased trial spacing enhances memory retrieval, independent of the impairing effect on memory encoding.

### The dmPFC is activated and necessary during the everyday memory task

We investigated whether the effects of trial spacing on memory strength were mediated by the dmPFC. To validate that training in the everyday memory task activates the dmPFC, we quantified neuronal activation during encoding using expression of the immediate early gene c-Fos (**Figure 2A**). c-Fos expression in the dmPFC was increased after training as compared to handled or home cage controls (**Figure 2B, C**). However, the number of c-Fos-expressing neurons was similar after training spaced with any encoding intertrial interval. This suggests that trial spacing did not increase the number of activated neurons during memory encoding.

**Figure 2.**
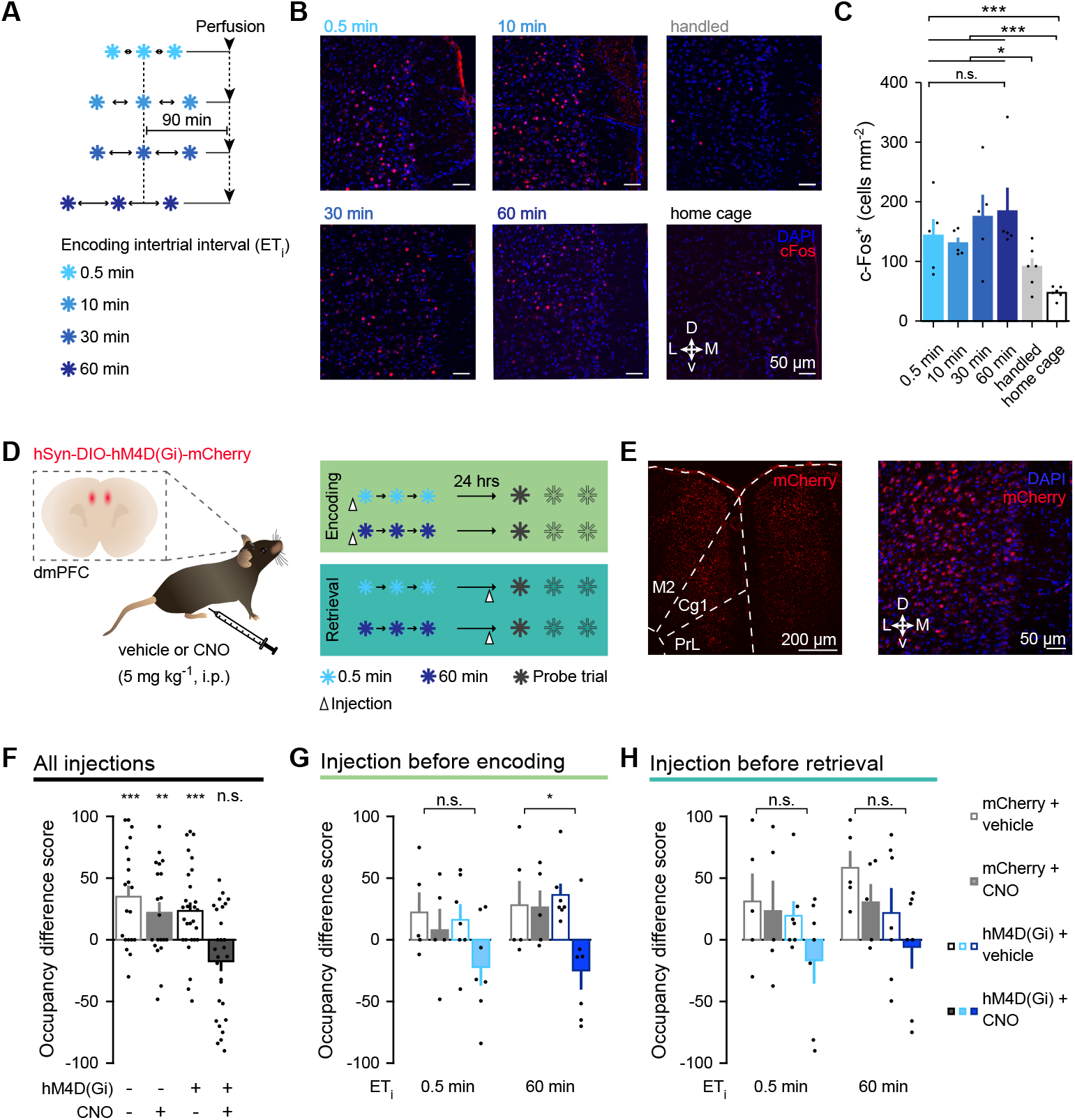
dmPFC activity is increased and necessary during the everyday memory task. **A** Timeline of behavioral procedures and tissue collection. **B** Representative images of c-Fos labeling in the dmPFC. **C** Training with any ET_i_ increased the number of cells expressing c-Fos as compared to home cage or handled controls (H_3_ = 15.8, p = 3.65·10^−4^; training vs. home cage: p = 7.71·10^−4^; training vs. handled: p = 0.013). Increasing the ET_i_ did not alter the number of c-Fos-expressing cells (H_4_ = 1.19, p = 0.754). **D** Chemogenetic silencing experiment. **E** Representative images of hM4D(Gi)-mCherry-expressing neurons in the dmPFC. **F** Memory on probe trials was impaired after silencing the dmPFC by injecting CNO into mice expressing hM4D(Gi)-mCherry (data pooled across injection time points and encoding intertrial intervals; mCherry + vehicle vs. chance: t_19_ = 3.87, p = 5.13·10^−4^; mCherry + CNO vs. chance: t_19_ = 2.56, p = 9.51·10^−3^; hM4D(Gi) + vehicle vs. chance: t_19_ = 3.45, p = 9.61·10^−4^; hM4D(Gi) + CNO vs. chance: t_19_ = −2.14, p = 0.979). **G** Memory was significantly reduced after silencing the dmPFC using CNO during spaced encoding (ET_i_ 60 min [right]; F_4,20_ = 4.16, p = 0.019, group, *t* = 1.46, p = 0.161; drug, *t* = 1.18, p = 0.251; interaction, *t* = −2.51, p = 0.021) but not massed encoding (ET_i_ 0.5 min [left]; F_4,20_ = 1.08, p = 0.179, group, *t* = 0.37, p = 0.718; drug, *t* = 0.19, p = 0.855; interaction, *t* = −0.77, p = 0.448). **H** Same as in (**G**), for retrieval (ET_i_ 0.5 min [left]; F_4,20_ = 1.31, p = 0.298, group, *t* = 0.27, p = 0.788; drug, *t* = 0.32, p = 0.753; interaction, *t* = −0.73, p = 0.473; ET_i_ 60 min [right]; F_4,20_ = 3.21, p = 0.062, group, *t* = −0.64, p = 0.526; drug, *t* = −0.47, p = 0.642; interaction, *t* = 0.01, p = 0.990). Cg1: cingulate cortex, area 1, M2: secondary motor cortex, PrL: prelimbic cortex. Scale bars 50 μm (B), 200 μm (E, left), 50 μm (E, right). Bars indicate mean (±SEM) across mice, black dots indicate data from a single mouse. * p < 0.05, ** p < 0.01, *** p < 0.001, n.s. non-significant.

To establish a causal role of the dmPFC, we chemogenetically inhibited it. We bilaterally transduced excitatory dmPFC neurons with the inhibitory chemogenetic tool hM4D(Gi), which is activated by clozapine- *N* -oxide (CNO; **Figure 2D, E**; ^34^). We first verified receptor function in dmPFC *ex vivo* electrophysiological recordings and established that CNO application to acute brain slices reduced the excitability of dmPFC neurons expressing hM4D(Gi) (**Figure S6**). Next, we addressed the role of dmPFC activity during the everyday memory task using a full factorial 2^4^ design (see Method Summary; **Figure 2D**). In well-trained mice expressing either hM4D(Gi) (n = 7) or mCherry (n = 5), we injected either vehicle or CNO at either of two time points (i.e. before memory encoding or before retrieval) using either of two encoding intertrial intervals (i.e. 0.5 min and 60 min; **Figure 2D**). Pooling data across time points and intervals showed that CNO-mediated inhibition impaired memory retrieval in hM4D(Gi)-expressing mice (**Figure 2F**). CNO application similarly reduced memory retrieval between individual time points and interval durations (**Figure 2G, H**). However, evaluating the individual time points and interval durations revealed that memory retrieval was only significantly influenced when the dmPFC was inhibited during spaced (i.e. 60 min) encoding. Overall, we conclude that memory storage requires dmPFC activation.

### Trial spacing increases the stability of the prefrontal cortex activation pattern

The major aim of this study was to evaluate whether trial spacing stabilizes the activity patterns of the neuronal populations throughout a session, i.e. whether it facilitates reactivation of a similar neuronal ensemble in subsequent trials. To this end, we used *in vivo* calcium imaging to simultaneously measure the activity patterns of on average 210 ± 99 (SD) individual dmPFC neurons per session in freely-moving mice (n = 499 sessions across 19 mice; **Figure 3A, B, S1E**; ^35^). After gaining optical access to the dmPFC with an implanted microprism, we used a miniaturized microscope (Doric Lenses) to image neurons expressing the calcium indicator GCaMP6m (**Figure 3C, Figure S7A–E**; ^36^). We ensured that carrying the miniaturized microscope did not hamper the mouse’s motility in the radial arm maze (**Figure S7F, G**). Using the constrained nonnegative matrix factorization for microendoscopic data (CNMF-E) algorithm ^37^, we extracted neuronal calcium activity and used the deconvolved inferred spike rate for further analysis (see Supplemental Methods). We computed the probability of a neuron being active by comparing the inferred spike rate in each trial to the pre-trial baseline period, using temporal subsampling to control for session duration (p_active_; see Supplemental Information; **Figure S8A–C**). We subsequently concatenated these values into an ensemble response vector and stacked the single-trial ensemble response vectors into an ensemble response matrix (n neurons × 6 trials; **Figure 3D**). The Pearson correlation between the rows of this matrix was used as the sessions trial-to-trial ensemble stability measure (**Figure 3D**).

**Figure 3.**
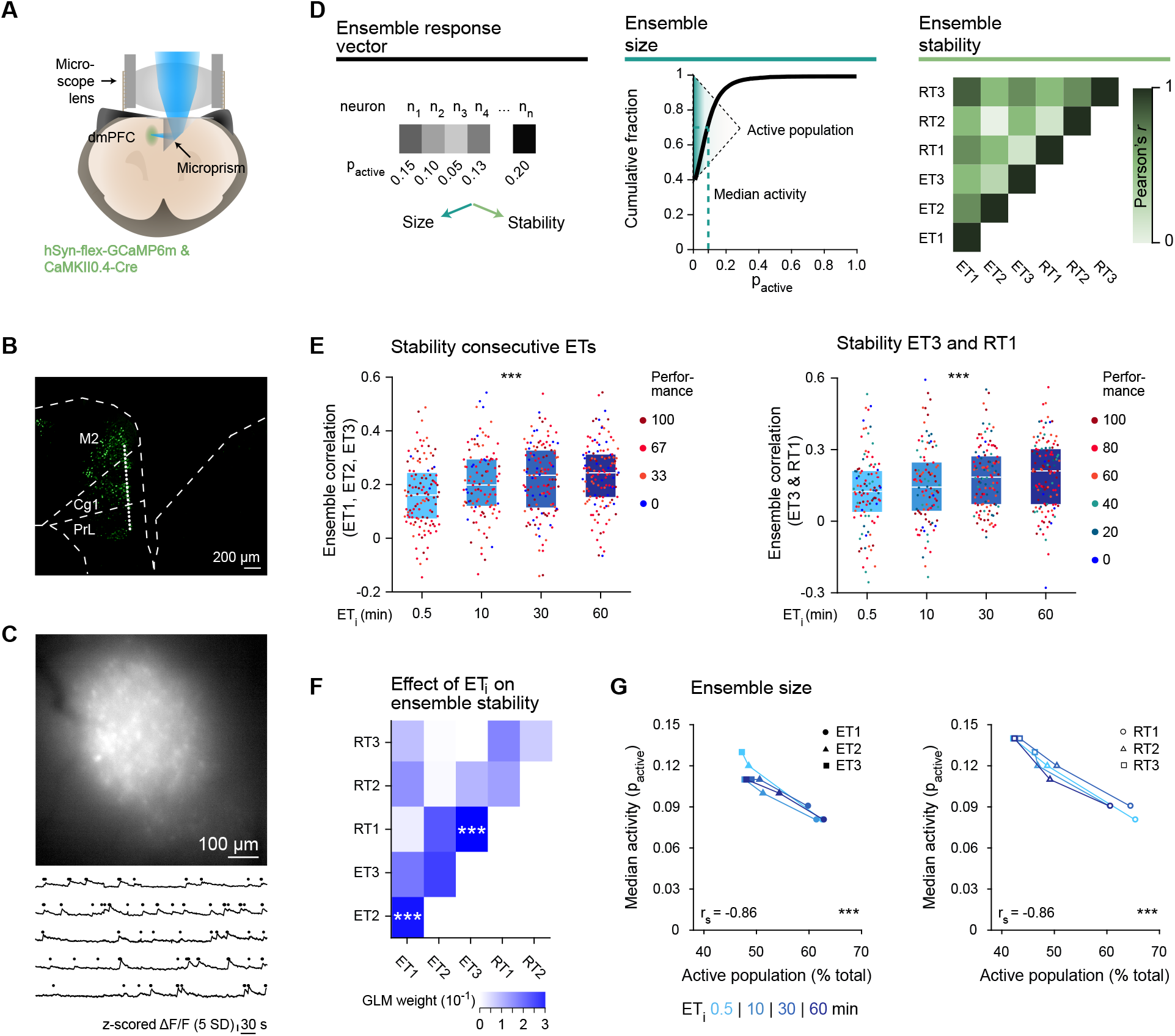
Trial spacing enhances ensemble stability, but not ensemble size. **A** Schematic of imaging preparation. **B** Approximate imaging plane (dotted line). **C** Sample frame showing dmPFC neurons (top), calcium activity and deconvolved spikes (traces and dots, bottom). **D** Schematic showing the quantification of ensemble size and stability. For each trial, the activity measure “p_active_” of individual neurons was concatenated into an ensemble response vector (left). Ensemble size: the active population (p_active_ > 0) and its median activity were inferred from the cumulative distribution of each ensemble response vector (middle). Ensemble stability: correlation between all ensemble response vectors of a session (right). **E** Ensemble correlation between consecutive encoding trials (ET1, ET2, and ET3; left) and between ET3 and RT1 (right) on individual sessions (n = 499) was higher for longer ET_i_s (consecutive encoding trials: F_4,495_ = 10.7, p = 2.89·10^−5^; ET_i_: *t* = 4.39, p = 1.71·10^−4^; ET3–RT1: F_4,495_ = 7.03, p = 9.93·10^−4^; ET_i_: *t* = 3.71, p = 2.37·10^−4^). Performance did not significantly covary with the ensemble correlation (consecutive encoding trials: *t* = 1.75, p = 0.08; ET3–RT1: *t* = 0.84, p = 0.40). **F** GLM weight of ET_i_ as predictor for ensemble correlation. The ensemble correlation between ET1 and ET2, as well as ET3 and RT1, depended on ET_i_ (ET1 vs. ET2: H3 = 17.5, p = 5.64·10^−4^; ET3 vs. RT1: H3 = 15.5, p = 1.45·10^−3^). **G** Both the size of the active population and its median activity depended on the trial identity (i.e. first, second or third trial) but not the ET_i_ (size active population F_4,495_ = 51.0, p = 1.85·10^−22^; trial identity: *t* = −10.1, p = 1.63·10^−23^; ET_i_: *t* = 0.49, p = 0.62; median activity of active population: F_4,495_ = 86.8, p = 2.83·10^−37^; trial identity: *t* = 13.0, p = 1.01·10^−37^; ET_i_: *t* = −1.98, p = 0.048). Cg1: cingulate cortex, area 1, M2: secondary motor cortex, PrL: prelimbic cortex, SD: standard deviation. Scale bars: 200 μm (B), 100 μm (C). *** p < 0.001.

The ensemble correlation between the first and second encoding trial was enhanced when the intertrial interval was longer, establishing that the ensemble reactivated more precisely (**Figure 3E, F**). Furthermore, trial spacing increased the ensemble correlation between the third encoding trial and the first retrieval trial, suggesting that the population activity pattern present during learning was more likely to be reactivated during retrieval (**Figure 3E, F**). The effect of trial spacing on ensemble correlation was not dependent on behavioral performance. As an alternative measure for similarity, we calculated the Euclidian distance between ensemble response vectors, which yielded similar results (**Figure S8D**). Overall, we find that increased trial spacing enhanced reactivation of the ensemble activity pattern instilled during encoding, while simultaneously strengthening memory retention.

### The size of the neuronal ensemble is not affected by trial spacing

We evaluated whether trial spacing altered the size of the neuronal ensemble (**Figure 3D**). From the cumulative distribution of each trial’s ensemble response vector, we inferred the active fraction within the neuronal population (i.e. the neuronal ensemble, p_active_ > 0) and the median activity of that population (**Figure 3D**). Interestingly, the neuronal ensembles became smaller across subsequent encoding and retrieval trials (i.e. first vs. second vs. third trial; **Figure 3G**). In addition, the median activity of the neuronal ensemble increased across subsequent trials (**Figure 3G**), indicating that the ensemble became sparser, yet the single neurons responded more strongly (see Discussion). However, neither ensemble size (i.e. the relative number of active neurons), nor its median activity was altered by trial spacing (**Figure 3G**).

With the overall ensemble size remaining stable, the memory enhancing effect of trial spacing could be attributed to a shift in the fraction of neurons preferentially responding to task-related events. We identified eight task-related behavioral variables that correlated with reward, motor activity, and decision-making: reward onset, reward approach (i.e. the final entry into the arm containing the rewarded sandwell), acceleration, speed, digging onset, digging offset, entry into the center platform, and intra-arm turns (**Figure S9A–C**; ^38^). On first inspection, neuronal responses did not appear time-locked or consistently occurring with the onset of these defined behaviors (**Figure S9D**). This was likely related to the naturalistic character of the everyday memory task, in which the individual components that comprise a behavior can occur simultaneously, whereas these appear discrete in more controlled experimental settings.

To determine whether the activity of individual neurons was modulated by task-relevant behaviors, we implemented an encoding model (generalized linear model, GLM). The model fitted the eight aforementioned behavioral variables as time-varying predictors of a neuron’s binarized inferred firing activity (**Figure 4A, B**, see Supplemental Information; ^39^). A neuron was classified as responsive to one of these behavioral variables if the weight of its corresponding time-varying predictor was significantly different from zero. Decoding performance was better upon training the encoding model with observed as compared to permuted inferred firing activity (**Figure 4C**). Across sessions, 22.7% of neurons were significantly modulated by at least one behavioral variable, most often reward onset, approach to reward, and digging onset (19.9%, 16.2%, and 18.1% of the population of modulated neurons, respectively; **Figure 4D**). This shows that dmPFC neuronal activity during the everyday memory task is modulated by specific behavioral variables.

**Figure 4.**
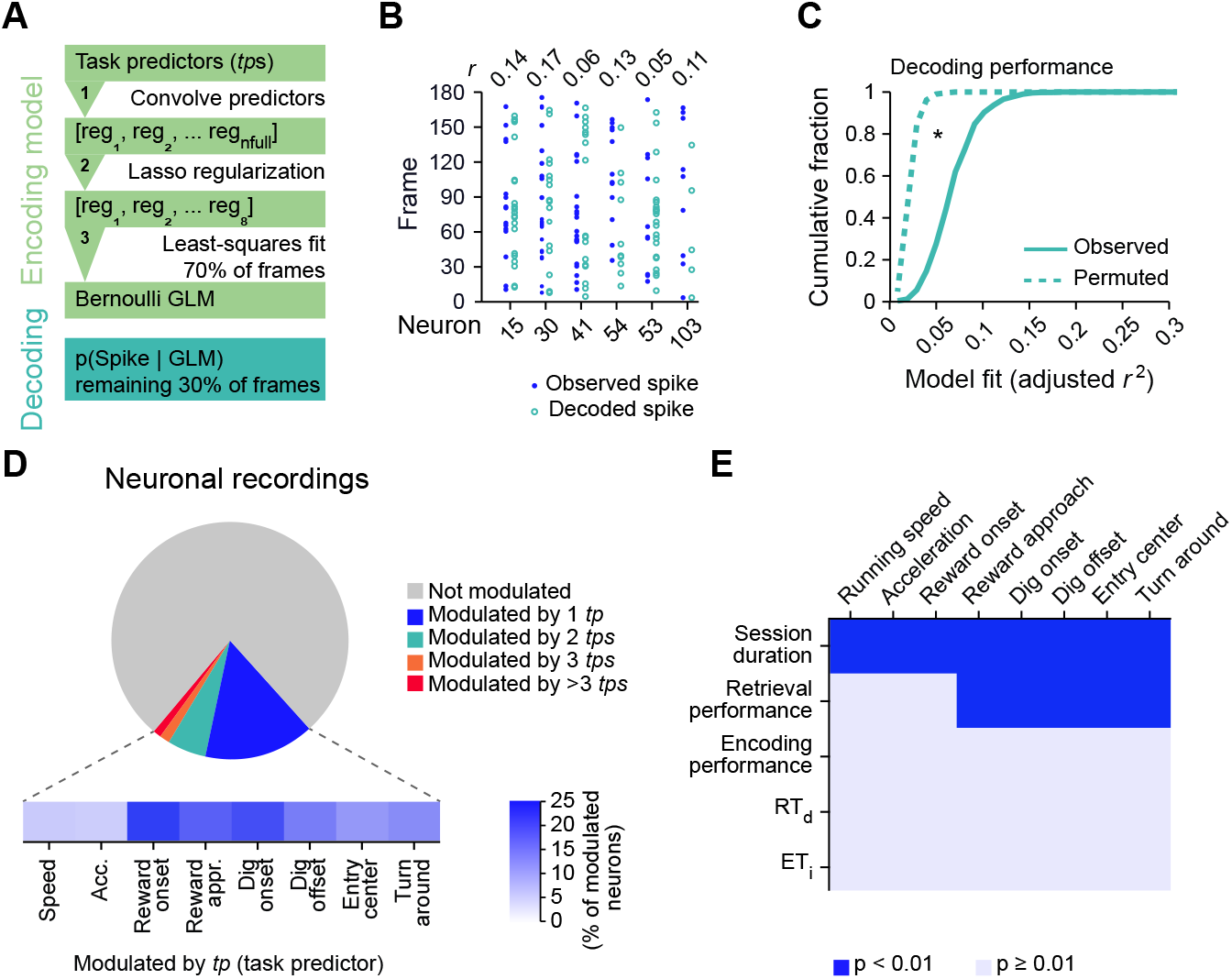
dmPFC neurons respond to multiple task-relevant behaviors irrespective of trial spacing. **A** Schematic of the generalized linear model. **B** Pearson’s *r* between observed and decoded spikes of the test subset for six randomly selected neurons. **C** Decoding performance was significantly better for observed than permuted spiking activity (observed vs. permuted: D_100_ = 0.21, p = 0.021). **D** Fractions of neurons (n = 105070 neurons from 499 sessions across 19 mice) that were modulated (top). Fractions of neurons modulated by the respective task predictor (bottom). **E** The effect of the five features session duration, performance on RT1 (“retrieval performance”), summed performance across ETs (“encoding performance”), ET_i_, and RT_d_, on the fraction of neurons modulated by the eight *tp*s. These five features differentially affected the fraction of neurons modulated by running speed (F-statistic vs. constant model [F]_5,494_ = 27.9, p = 6.95·10^−25^; session duration [SeD]: *t* = 9.80, p = 7.65·10^−21^), acceleration (F_5,494_ = 5.21, p = 1.13·10^−4^; SeD: *t* = 3.25, p = 1.22·10^−3^), reward onset (F_5,494_ = 28.6, p = 1.62·10^−25^; SeD: *t* = 9.84, p = 5.70·10^−21^), reward approach (F_5,494_ = 31.1, p = 1.55·10^−27^; SeD: *t* = 9.11, p = 2.04·10^−18^, retrieval performance [RP]: *t* = 4.27, p = 2.35·10^−5^), dig onset (F_5,494_ = 38.9, p = 1.05·10^−33^; SeD: *t* = 11.2, p = 4.22·10^−26^, RP: *t* = 3.51, p = 4.86·10^−4^), dig offset (F_5,494_ = 38.9, p = 1.97·10^−33^; SeD: *t* = 10.9, p = 3.65·10^−25^, RP: *t* = 2.95, p = 3.30·10^−3^), entry into the center platform (F_5,494_ = 39.2, p = 6.21·10^−34^; SeD: *t* = 10.5, p = 2.38·10^−23^, RP: *t* = 4.68, p = 3.64·10^−6^), and intra-arm turns (F_5,494_ = 25.9, p = 2.80·10^−23^; SeD: *t* = 7.97, p = 1.07·10^−14^, RP: *t* = 3.77, p = 1.82·10^−4^). All post hoc tests were Bonferroni corrected for five comparisons, only significant post hoc tests are reported. Acc: acceleration, appr: approach, GLM: generalized linear model, *tp*: task predictor, reg: regressor. * p < 0.05.

However, trial spacing did not have a significant influence on the fractions of behaviorally modulated neurons, nor did the encoding performance or retrieval delay duration (**Figure 4E**). Firing modulation by all identified task-relevant behaviors did depend on session duration (**Figure 4E**), likely because an increased session duration inherently produces more neuronal spikes and therefore data for the GLM to fit. Furthermore, retrieval performance correlated with the fraction of neurons modulated by certain behavioral variables, reward approach, dig onset, dig offset, entry into the center platform, and intra-arm turns. Therefore, we conclude that a sparse population of dmPFC neurons encoded task-related behaviors similarly across experimental conditions and we did not find evidence of an effect of trial spacing on the number of neurons involved. Overall, trial spacing in the everyday memory task enhances ensemble stability but it does not affect ensemble size.

## Discussion

We explored whether trial spacing strengthens memory by altering characteristics of the neuronal ensemble. We observed the behavioral effect of trial spacing on the everyday memory task and characterized the activity of prefrontal neurons that were necessary for task performance. During learning and upon memory retrieval, the ensemble activity pattern reactivated more precisely when trial spacing was increased. In contrast, trial spacing did not affect the overall size of the activated ensemble, nor the size of the subpopulations of neurons that responded to specific task-related behaviors. Our results suggest that more precise reactivation of the neuronal ensemble during spaced training strengthens connectivity that is conducive to memory retention and retrieval.

### Spaced training strengthens memory

Spaced training in the everyday memory task strengthens memory in rats ^7^ and we report the same in mice. Earlier studies investigating the effect of trial spacing on episodic-like memory in mice ^6^ and rats ^40^ have reported an inverted U-shaped relation, although the exact width and amplitude of the effect varied. Our study likewise reports an inverted U-shaped relation, as spacing trials at intervals of 60 min resulted in the strongest next-day (24 hrs) memory, while shorter (10 min and 30 min) and longer intervals (180 min) resulted in substantially poorer memory. The observed temporal window aligns with expectations from facilitated molecular signaling and synaptic physiology underlying the spacing effect ^2; 6; 8^.

As compared to spaced training, massed training in the everyday memory task affected memory in a rather complex manner. As expected, memory retrieval was poorer after massed training than after any spaced training regimen. Surprisingly, the ability to retrieve memory following massed training was better after 24 hrs, and even 48 hrs, as compared to 2.5 hrs. We propose that memory acquired during massed training might have only been stabilized after several hours. A similar phenomenon has been reported during massed motor learning in mice, in which both memory stabilization and concomitant synaptic remodeling occurred delayed as compared to spaced motor learning ^5^. However, delayed memory stabilization was not observed in two earlier studies using the everyday memory task ^7; 41^. This variation can possibly be attributed to methodological differences such as the animal model ^7; 41^, the number of encoding trials ^41^, navigational strategy ^33^, handling, or intertrial sleep epochs.

### Prefrontal activity in the everyday memory task

We focused our neuronal recordings and manipulations on the dmPFC. Activity of dmPFC neurons correlated with a range of task-relevant events on the everyday memory task, most notably reward (anticipation) and motor behavior, which is consistent with other reports in rodent PFC ^38; 42^. We established a causal link between prefrontal activity and memory formation by chemogenetically inactivating the dmPFC, which impaired next-day memory. This seemingly conflicts with reports that inactivation of prefrontal areas disrupts remote but not recent memories ^43^. However, the early dependence of task-instilled memories on the dmPFC may have followed from accelerated systems consolidation, as observed in other behavioral paradigms where learning occurred within the context of relevant pre-existing knowledge ^30^.

Episodic-like memories formed in the everyday memory task unlikely depended solely on the dmPFC. Specifically the hippocampus ^44^ and retrosplenial cortex ^45^ have long been implicated in various forms of declarative memory. Indeed, the hippocampus and prefrontal cortex have been suggested to perform complementary roles in episodic-like memory processing ^31; 46^. Furthermore, retrosplenial neurons form ensembles that stabilize during learning of spatial reference memory tasks ^47^ and the stability of these retrosplenial ensembles can predict memory retention ^48^. Interestingly, a recent study shows that trial spacing upregulates a variety of genes, including immediate early genes, in both hippocampus and retrosplenial cortex in the rat ^7^. Whether and how neuronal ensembles in the mouse hippocampus and retrosplenial cortex are affected by spaced training in the everyday memory task would be of interest for future investigation.

### The spacing effect, synaptic strength, and memory stability

Our experiments explored the possibility that trial spacing enhances memory by altering the size or stability of a neuronal ensemble. We quantified ensemble size using two distinct methods. First, we determined the neuronal ensemble size using calcium imaging of GCaMP6-expressing neurons, which closely reflects the temporal dynamics of neuronal firing throughout each trial ^36^. This approach allowed for detecting both highly active and transiently activated neurons, while controlling for the influence of training duration on ensemble size by temporal subsampling. Second, we quantified ensemble size from the number of c-Fos-expressing neurons after a full encoding session. This method is more likely to only include strongly activated neurons that subsequently underwent plasticity implicated in long-term memory storage ^49^. Despite the methodological differences between these approaches, both yielded similar results: the size of the active population was not influenced by trial spacing. This is in agreement with the previous observation that ensemble size is generally quite stable and is not strongly influenced by factors such as the type of memory and the strength of a memory ^24^.

Irrespective of trial spacing, behavioral training activated a progressively smaller population of neurons, whose activity was stronger than in previous trials. Sparsening of the neuronal ensemble can enhance memory selectivity, as for instance observed during *Drosophila* olfactory conditioning ^26^. Several studies propose that this is the consequence of a competitive process ^9; 24^, in which highly excitable pyramidal neurons exclude less excitable neighboring pyramidal neurons from becoming part of the neuronal ensemble via local inhibition. A similar process might ensure ensemble sparsity in the everyday memory task, thereby balancing memory fidelity with memory capacity ^9^.

The main consequence of trial spacing was that the neuronal ensemble reactivated in a pattern more reminiscent of previous learning experiences, corroborating theoretical predictions ^50^ and reports in human subjects ^23^. We suggest that more precise ensemble reactivation reflects specific synaptic processes that underlie memory formation ^14^. One such process is CaMKII activation, which unfolds on a similar timescale as spacing-induced memory enhancement and has previously been implicated in the spacing effect ^2^. A major outstanding question is whether these synaptic processes affect a random population of synapses or are confined to previously tagged synapses, as predicted by the synaptic tagging and capture theory ^51^. This could be addressed using *in vivo* imaging of structure and function of individual spines during the everyday memory assay ^52^.

Overall, our data show that trial spacing increases the strength of connectivity within the ensemble, supposedly making memory more robust and increasing the probability of memory retrieval. Our findings provide the first direct description of how activity of the same neuronal population during memory encoding and retrieval mediates the spacing effect, a phenomenon originally described over a century ago ^1^.

## Acknowledgments

We thank Claudia Huber, Andreas Kucher, Max Sperling, Volker Staiger, Helena Tultschin, and Frank Voss for technical assistance; Sandra Reinert for help with the implant method and discussions; Julia Kuhl for illustrations. This project was funded by the Max Planck Society and the Collaborative Research Center SFB870 of the German Research Foundation (DFG) to T. B. and M. H.

## Author Contributions

All authors designed the study. A. G. acquired the data. A. G. and P. G. analyzed the data. T. B. and M. H. acquired funding. All authors wrote the manuscript.

## Declaration of Interests

The authors have declared that no competing interests exist.

## Method Summary

For greater detail, refer to Supplemental Methods.

### Mice

Female C57BL/6NRj mice (∼ postnatal day 90 at experimental onset) were communally housed in enriched, individually ventilated cages. All procedures were performed in accordance with the institutional guidelines of the Max Planck Society and the local government (Regierung von Oberbayern, Germany).

### Surgical procedures

Mice were anesthetized with a mixture of fentanyl, midazolam, and medetomidine in saline (0.05 mg kg^−1^, 5 mg kg^−1^, and 0.5 mg kg^−1^ respectively, injected intraperitoneally). A head plate implantation was carried out as previously described ^53^. For imaging experiments, a 3 mm circular craniotomy was created (centered at AP 2.0 mm, ML 0.75 mm from bregma) and a viral vector mixture of AAV2/1:CamKII0.4-Cre (4.6·10^9^ GC ml^−1^), and AAV2/1:hSyn-flex-GCaMP6m (3.2·10^12^ GC ml^−1^) was unilaterally injected into the dmPFC (150 nl injection^−1^). Subsequently, a microprism was implanted by removing the dura over one hemisphere and lowering the microprism into the sagittal fissure, facing the other hemisphere.

For chemogenetic inactivation experiments, a viral vector mixture of AAV2/1:CamKII0.4-Cre (2.1·10^11^ GC ml^−1^), and either AAV2/9:hSyn-DIO-mCherry (2.1·10^12^ GC ml^−1^) or AAV2/9:hSyn-DIO-hM4D(Gi)-mCherry (2.3·10^12^ GC ml^−1^) was bilaterally injected into the dmPFC (150 nl injection^−1^). After surgery, anesthetic agents were antagonized with a mixture of naloxone, flumazenil, and atipamezole in saline (1.2 mg kg^−1^, 0.5 mg kg^−1^, and 2.5 mg kg^−1^ respectively, injected subcutaneously [s.c.]). Mice received carprofen (5 mg kg^−1^, injected s.c.) and dexamethasone (2 μg kg^−1^, injected s.c.) for two subsequent days.

### Behavioral procedures

Mouse handling, training, and testing was performed similarly as previously described ^7^, with the main exception that training was conducted in a custom-made radial arm maze (**Figure 1A, S1A–C**). Behavioral training was recorded with an overhead video camera and the frame-by-frame position of the mouse was automatically annotated using custom-written MATLAB software. Each session typically consisted of three encoding trials (ETs) and three retrieval trials (RTs; **Figure 1B**). The ETs had an intertrial interval of either 30 s (“massed”), 10 min, 30 min, or 60 min (all “spaced”) and the delay between the final ET and first RT was 2.5 hrs or 24 hrs (**Figure 1B**). For each ET and RT, the number of erroneous sandwell digs was counted. For each trial, the performance index (PI) was calculated as

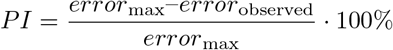

with error_max_ being 1 for ETs and 5 for RTs. Probe trial sessions were conducted as normal, however the rewarded sandwell did not contain a reward for the first 60 s of the first retrieval trial. Performance on probe trials was quantified as the relative dig time at the rewarded sandwell (“occupancy difference score”). For chemogenetic inactivation, mice were injected intraperitoneally with either vehicle or CNO (5 mg kg^−1^) 45 min before behavioral testing (**Figure 2D**).

### Histology and immunohistochemistry

For experiments quantifying immediate early gene expression, mice were perfused 90 min after the onset of the second ET (**Figure 2A**). Brains were sectioned on a microtome (40 μm, coronal) and every 5^th^ section containing dmPFC was stained for c-Fos and DAPI. Micrographs containing the dmPFC were acquired using a laser-scanning confocal microscope (TCS SP8, 20× NA 0.75 objective) and analyzed by counting the number of c-Fos immunopositive neurons in a blind manner in the dmPFC.

### Miniaturized microscopy and data processing

Images were acquired with a commercially available miniaturized microscope (Basic Fluorescence Microscopy System - Surface, Doric Lenses) at a frame rate of 10 Hz and a resolution of 630 × 630 pixels (field-of-view 1 mm^2^). Laser power under the objective lens (2× magnification, 0.5 NA) was <1 mW for all imaging experiments. Image registration, motion correction, intrasession frame concatenation, and source extraction were carried out using custom implementations of the NoRMCorre and CNMF-E algorithms ^37; 54^. We used a probabilistic measure (p_active_) to quantify the activity of an individual neuron during an individual trial, controlling for trial length (**Figure S8C**).

### Generalized linear model (GLM) quantifying behavioral modulation of neuronal responses

A GLM was fitted to the spiking activity of single neurons to establish the predictive power of relevant task predictors (*tp*s) for neuronal activity (**Figure 4A**). Briefly, a design matrix containing the *tp*s and the spiking dataset (70% of session’s frames) were supplied to the MATLAB function “*fitglm*”. When a resulting regression coefficient was significant after Bonferroni correction, the neuron was classified as modulated by this task predictor. The model’s decoding performance was quantified by correlating the observed spiking responses (remaining 30% of session’s frames) with the responses predicted by the GLM using the MATLAB function “*predict*”.

### Statistical analysis

All data are presented as mean (± SEM) unless stated otherwise. Normality of distributions was assessed using the Kolmogorov-Smirnov test and appropriate parametric or non-parametric tests were used. Parametric analyses included the Student’s t-test (test statistic t) and general linear models including one-way ANOVA (test statistic F) for data consisting of two groups or more than two groups, respectively. Non-parametric analyses for data consisting of two groups included the Kolmogorov-Smirnov test (test statistic D), the Mann-Whitney U test (test statistic U), Spearman correlation (test indicated by Spearman’s rank correlation coefficient r_s_), and the Wilcoxon’s rank-sum test (test statistic W). Non-parametric analyses for data consisting of more than two groups included Pearson’s chi-square test (test statistic X^2^) and the Kruskall-Wallis test (test statistic H). For all statistical tests, alpha was set at 0.05 and tests were conducted two-tailed unless stated otherwise. In case of multiple comparisons, a Bonferroni alpha correction was applied.

## Notes

https://figshare.com/s/82bcda77a0435f199438

https://figshare.com/s/b90b7171036bc3849f9c

https://figshare.com/s/c3a41dfc9cdfcdda0ffb

### Competing Interest Statement

The authors have declared no competing interest.

