## Supplemental Information for "Spaced training enhances memory and prefrontal ensemble stability in mice"

Supplemental Information includes:

Supplemental Figures 1-9
Supplemental Table 1
Supplemental Methods

### Supplemental Figures

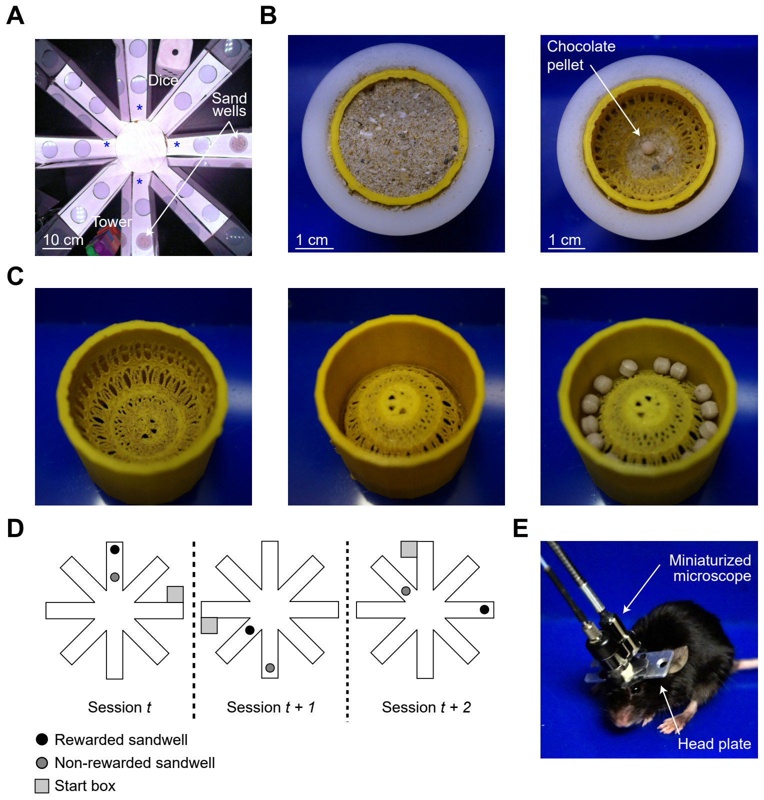


**Supplemental Figure 1. Experimental setup for behavioral testing and miniature microscope imaging.** **A** Overhead photograph of the 8-arm radial maze with two proximal landmarks (LEGO® DUPLO® tower and plush dice). Each arm contained two circular openings that could be covered with Plexiglas lids or contain a sandwell. Blue asterisks mark cardinal arms. **B** Overhead photograph of the sandwell filled with sand (left) and without sand (right), revealing the chocolate flavored food pellet. **C** Overhead photographs of the sandwell inlay, oriented top-up (left) or top-down (middle). The inlay was filled with chocolate flavored food pellets (right) to provide the same olfactory cues for all sandwells. **D** Example maze layouts on three consecutive sessions. Note that the location of the start box, the rewarded sandwell and non-rewarded sandwell were altered between sessions. **E** Mouse carrying a miniaturized microscope, mounted on a head plate implant.

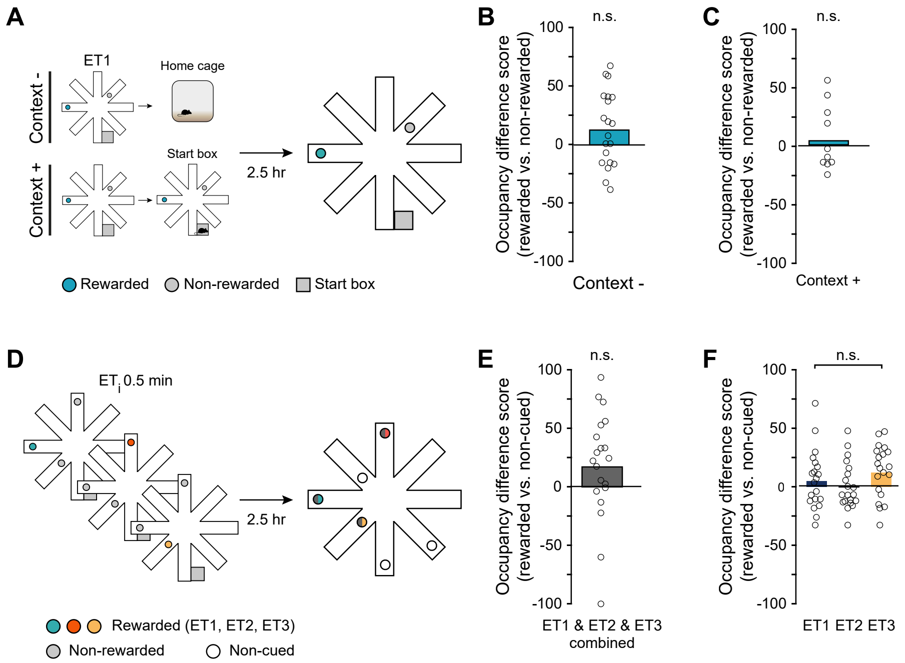


**Supplemental Figure 2. Only congruent, multi-trial training resulted in memory on the everyday memory task. A** Single trial learning experiment. A single encoding trial was conducted, after which the mouse was kept in the home cage during the entire retrieval delay (no re-introduction: Context -), or briefly re-introduced into the start box (Context +). After a retrieval delay of 2.5 hrs, a probe trial was conducted, and the occupancy at the rewarded and non-rewarded sandwell was recorded. **B** Memory retrieval (occupancy difference) was not observed in the probe trial upon training with a single encoding trial without re-introduction (Context -: $W_{19} = 156$, $p = 0.057$, $n = 20$ mice). **C** Same as in (**B**) for a single encoding trial with re-introduction (Context +: $t_9 = 0.426$, $p = 0.679$, $n = 10$ mice). **D** Incongruent learning experiment. On ET1, ET2, and ET3 (encoding intertrial interval [$ET_i$] 0.5 min), the location of the rewarded sandwell was altered. After a retrieval delay of 2.5 hrs, a probe trial was conducted. **E** Occupancy was similar for previously rewarded sandwells and all non-cued sandwells (rewarded vs. non-cued sandwells: $W_{19} = 155$, $p = 0.062$, $n = 20$ mice) **F** Occupancy was similar for all of the previously rewarded sandwells (ET1 vs. ET2 vs. ET3: $H_2 = 2.03$, $p = 0.363$). Circles indicate data from each mouse and bars indicate population mean. n.s., non-significant.

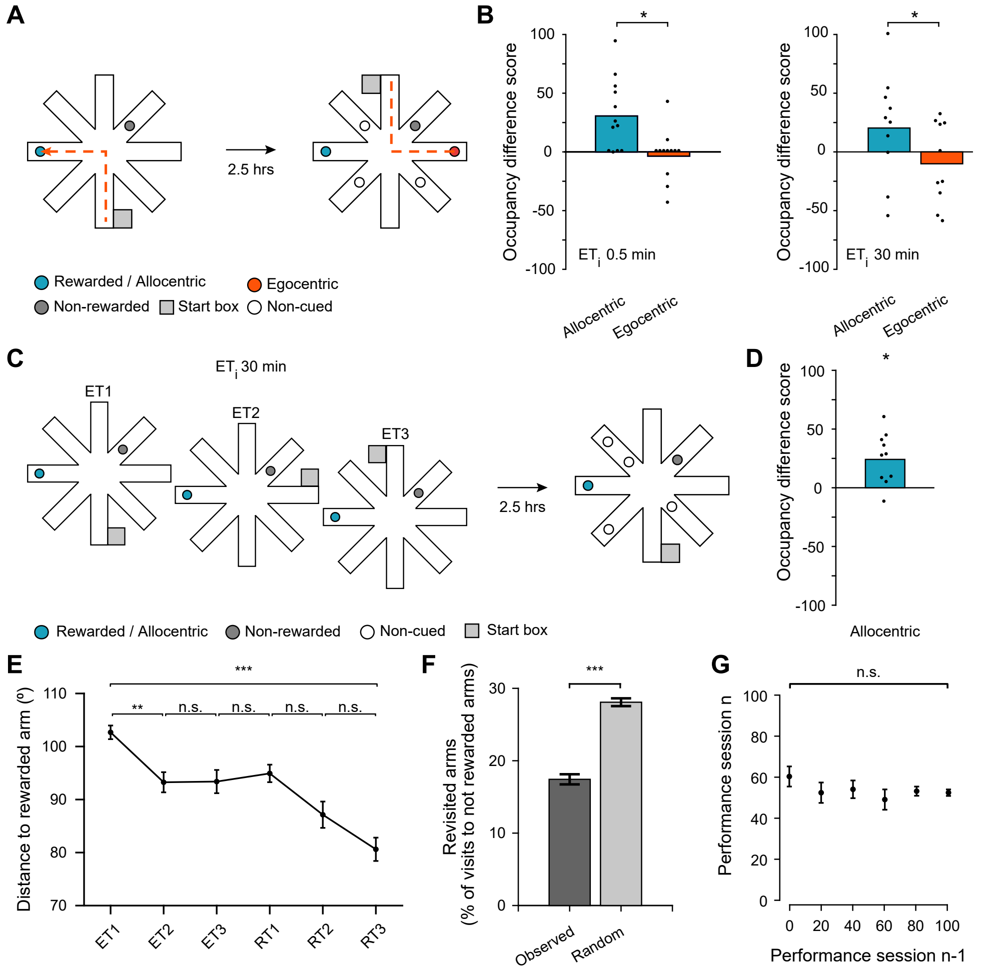


**Supplemental Figure 3. Navigational strategies on the everyday memory task.** **A** Navigational strategy experiment. Mice were trained on three encoding trials using either massed ($ET_i$ 0.5 min) or spaced ($ET_i$ 30 min) training. During the retrieval delay, the start box was placed at the opposite end of the maze, altering the path but not the location of the rewarded sandwell ("allocentric sandwell"). Egocentric spatial navigation would lead the mouse to a non-cued sandwell ("egocentric sandwell"). **B** The occupancy at the allocentric versus egocentric sandwell upon both massed (left: $t_{11} = 3.69$, $p = 3.59 \cdot 10^{-3}$, n = 12 mice) or upon spaced training (right: $t_9 = 2.40$, $p = 0.040$, n = 10 mice). **C** Forced allocentric strategy experiment. The start box location was altered after each encoding trial ($ET_i$ 30 min), enforcing allocentric navigation. **D** Occupancy at the rewarded sandwell (observed vs. chance [occupancy difference 0]: $t_9 = 3.50$, $p = 0.007$, n = 10 mice). **E** The mean angular distance of the arm the mouse was in, relative to the rewarded arm, decreased across trials ($H_5 = 67.4$, $p = 3.56 \cdot 10^{-13}$; ET1 vs ET3: $p = 1.69 \cdot 10^{-3}$). **F** The observed, relative fraction of incorrect arm visits was lower than expected from chance (observed vs. random arm visits; $t_{781} = -27.7$, $p = 2.59 \cdot 10^{-118}$). **G** The performance in RT1 of the previous session did not affect that of the current session ($H_5 = 3.97$, $p = 0.554$, n = 609 trials in 20 mice). Filled dots indicate data from each mouse, circles indicate the population mean ± SEM, bars indicate population mean, error bars indicate SEM. n.s., non-significant, * $p < 0.05$, ** $p < 0.01$, *** $p < 0.001$.

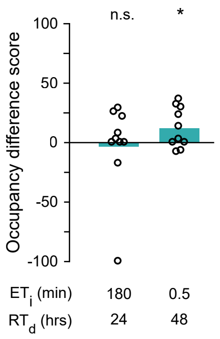


**Supplemental Figure 4. Extending encoding and retrieval spacing.** Occupancy at the rewarded sandwell upon training with an extended encoding intertrial interval ($ET_i$ 180 min, $RT_d$ 24 hrs; observed vs. chance [occupancy difference 0]: $U = 22$, $p = 0.641$, $n = 10$ mice) or an extended retrieval delay ($ET_i$ 0.5 min, $RT_d$ 48 hrs; observed vs. chance [occupancy difference 0]: $U = 40$, $p = 0.039$, $n = 10$ mice). Open circles indicate data from one mouse, n.s., non-significant, * $p < 0.05$.

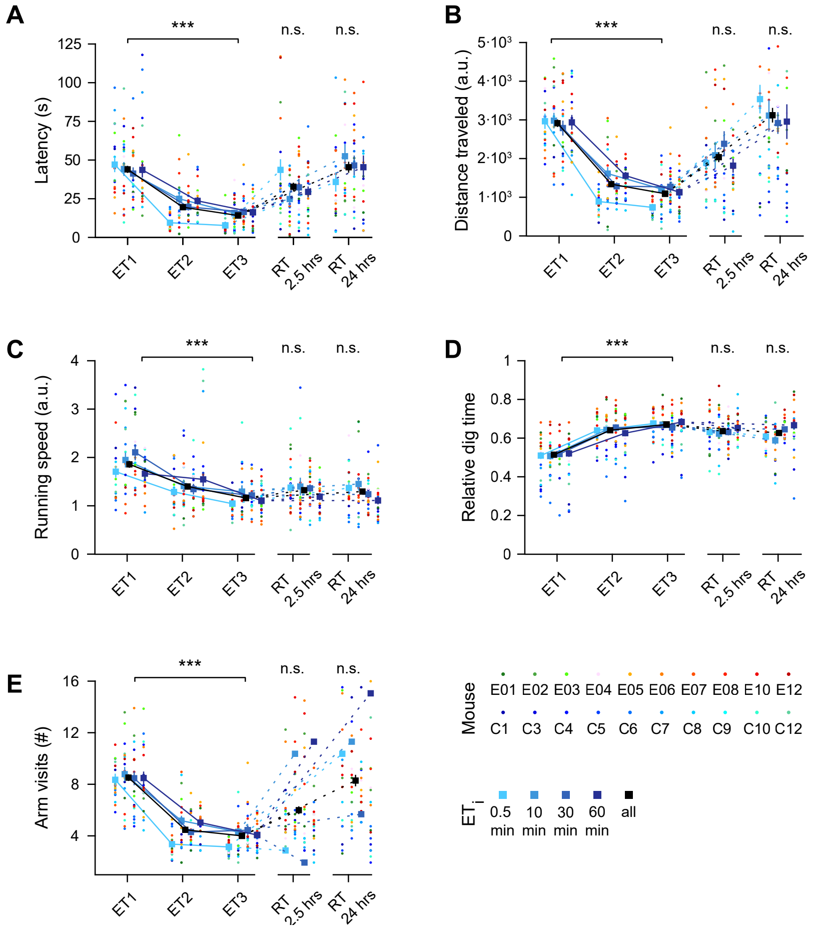


**Supplemental Figure 5. Behavioral parameters of individual mice in the everyday memory task.** The mean values of individual mice (n = 19) are represented by colored dots and are sorted by the session's $ET_i$ (0.5, 10, 30, or 60 min). The means for each trial, pooled across mice and $ET_i$s, are represented by black lines and squares. **A** Latency to the rewarded arm on ET1, ET2, and ET3 ("consecutive encoding trials", F-statistic vs. constant model [F] = 710, p = $1.96 \cdot 10^{-24}$; encoding trial identity [ETID]: $t$ = -11.7, p = $4.42 \cdot 10^{-25}$; $ET_i$: $t$ = 1.77, p = 0.078) and the first retrieval trial (RT1) after a retrieval delay ($RT_d$) of 2.5 hrs or 24 hrs (2.5 hrs: $r_s$ = -0.06, p = 0.622; 24 hrs: $r_s$ = 0.014, p = 0.909). **B** Same as in **A**, for the distance traveled (consecutive encoding trials: $F_{2,27}$ = 47.6, p = $6.27 \cdot 10^{-18}$; ETID: $t$ = 9.67, p = $1.05 \cdot 10^{-18}$; $ET_i$: $t$ = -1.24, p = 0.214; $RT_d$ 2.5 hrs: $r_s$ = -0.09, p = 0.456; $RT_d$ 24 hrs: $r_s$ = 0.221, p = 0.058). **C** Same as in **A**, for the running speed (consecutive encoding trials: $F_{2,27}$ = 111, p = $3.32 \cdot 10^{-34}$; ETID: $t$ = -14.8, p = $4.95 \cdot 10^{-35}$; $ET_i$: $t$ = 1.65, p = 0.101; $RT_d$ 2.5 hrs: $r_s$ = -0.07, p = 0.578; $RT_d$ 24 hrs: $r_s$ = -0.123, p = 0.297). **D** Same as in **A**, for the relative dig time (consecutive encoding trials: $F_{2,27}$ = 30.5, p = $2.01 \cdot 10^{-12}$; ETID: $t$ = -7.68, p = $4.97 \cdot 10^{-13}$; $ET_i$: $t$ = 1.40, p = 0.163; $RT_d$ 2.5 hrs: $r_s$ = -0.12, p = 0.328; $RT_d$ 24 hrs: $r_s$ = -0.21, p = 0.074). **E** Same as in **A**, for the number of arm visits (consecutive encoding trials: $F_{2,27}$ = 94.7, p = $1.81 \cdot 10^{-30}$; ETID: $t$ = -13.8, p = $1.54 \cdot 10^{-31}$; $ET_i$: $t$ = 0.55, p = 0.582; $RT_d$ 2.5 hrs: $r_s$ = -0.09, p = 0.426; $RT_d$ 24 hrs: $r_s$ = -0.11, p = 0.367). Squares indicate mean ± SEM across animals. n.s., non-significant, *** p < 0.001.

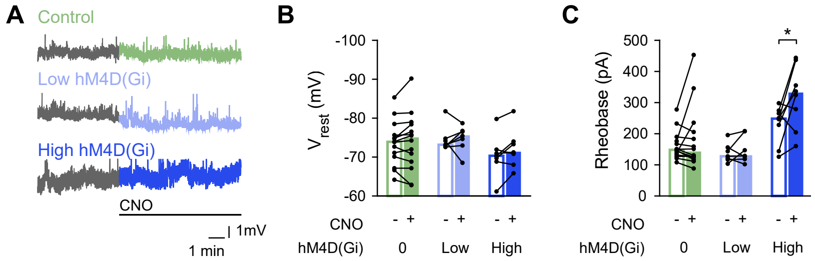


**Supplemental Figure 6.** ***Ex vivo*** **characterization of the effect of clozapine-*N*-oxide on hM4D(Gi)-expressing neurons. A** Example current-clamp recordings of the membrane potential of neurons not transduced (control; n = 14 neurons), transduced with a low titer (titer $2.3 \cdot 10^{10}$ GC $ml^{-1}$; n = 7 neurons), or transduced with a high titer (titer $2.3 \cdot 10^{12}$ GC $ml^{-1}$; n = 9 neurons) hM4D(Gi)-encoding AAV before and after clozapine-*N*-oxide (CNO) application. **B** CNO application did not affect the resting membrane potential ($V_{rest}$; $F_{2,52} = 1.06$, $p = 0.375$). **C** CNO application affected rheobase ($F_{2,54} = 10.5$, $p = 1.53 \cdot 10^{-5}$), post hoc analysis revealed an effect for high hM4D(Gi)-expressing neurons ($t_7 = -2.62$, $p = 0.034$). Recordings from an individual neuron (circles) are connected with lines, bars indicate population means. n.s., non-significant, * $p < 0.05$.

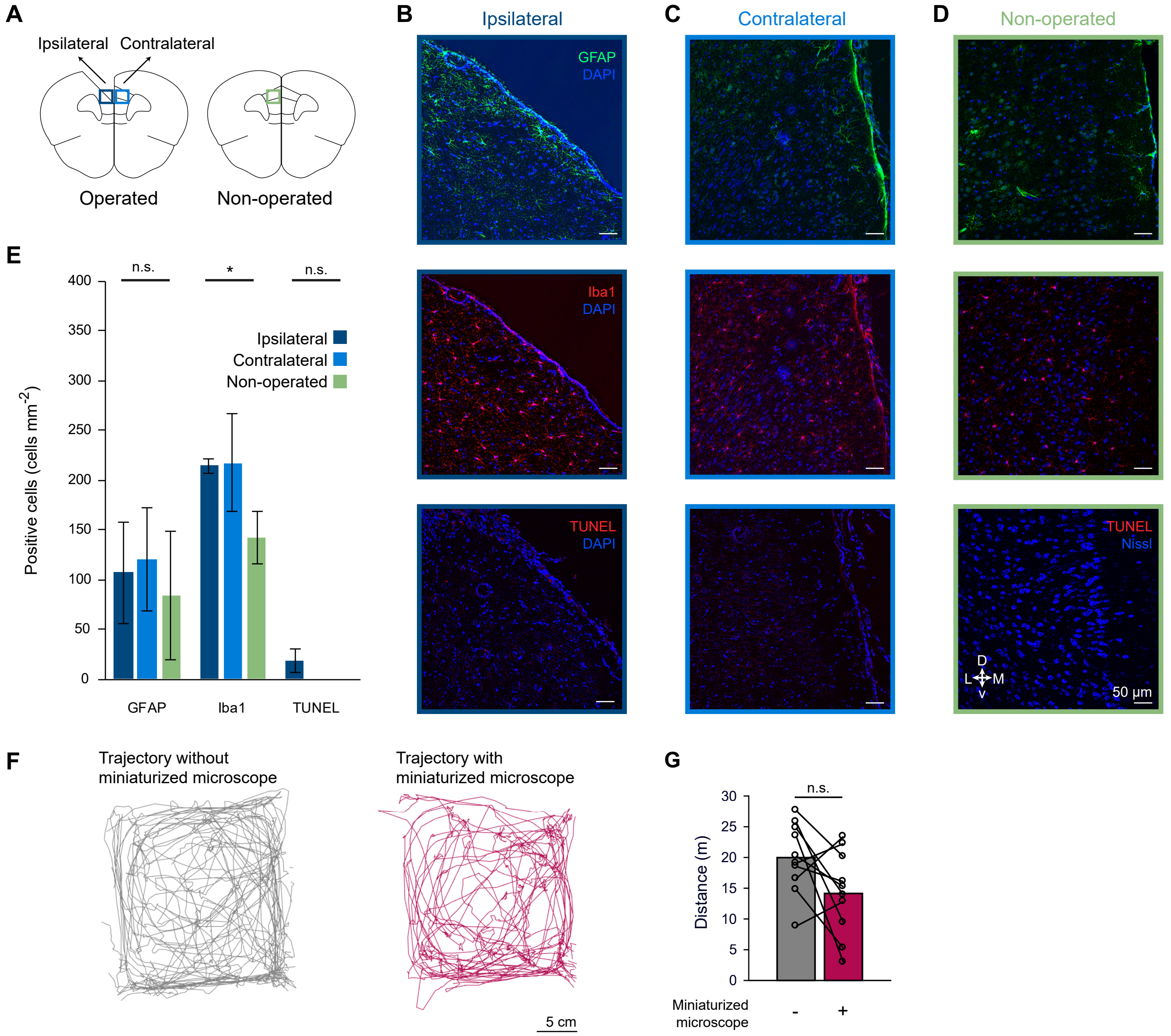


**Supplemental Figure 7. Control experiments regarding miniaturized microscope imaging of the dmPFC. A** A subsection of the ipsilateral and contralateral dmPFC hemisphere of mice with (operated) or without (non-operated) a microprism implant was analyzed for markers of astrogliosis (GFAP expression), microgliosis (Iba1 expression) and apoptosis (TUNEL assay). **B–D** Representative micrographs of dmPFC sections from the ipsilateral hemisphere (**B**), contralateral hemisphere (**C**), and non-operated hemisphere (**D**) labeled with GFAP (top), Iba1 (middle), and TUNEL (bottom). **E** The number of GFAP-expressing ($H_2$ = 0.35, p = 0.840), Iba1-expressing ($H_2$ = 6.05, p = 0.049), and TUNEL-positive ($H_2$= 3.85, p = 0.146) cells in the ipsilateral (n = 4 mice), contralateral (n = 4 mice) and non-operated (n = 3 mice) hemispheres. **F** Representative trajectories during a five-minute home cage exploration while mice did (right) or did not (left) carry the miniaturized microscope. **G** The distance mice traveled was not significantly affected by carrying the miniaturized microscope (- vs. +: T = 45, p = 0.084, n = 10 mice). Open circles indicate data from one mouse, bars indicate means. Scale bars 50 µm (**B–D**), 5 cm (**F**). n.s., non-significant, * $p < 0.05$.

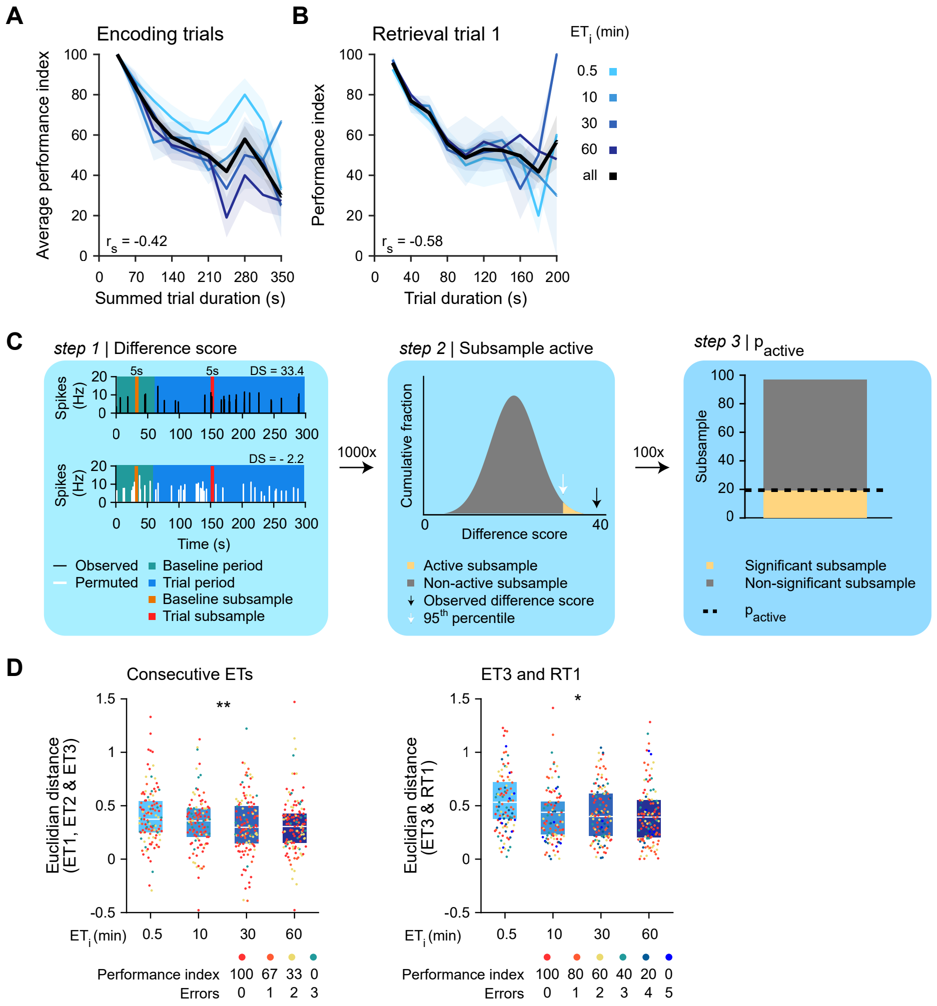


**Supplemental Figure 8. Quantification of the activity and stability of a neuronal population.** **A** The behavioral performance on, and duration of, all encoding trials of a session, sorted by $ET_i$ (n = 499 sessions; $F_{4,495} = 47.7$, $p = 1.12{\cdot}10^{-19}$; performance: $t = -9.67$, $p = 2.18{\cdot}10^{-20}$; $ET_i$: $t = -0.49$, $p = 0.623$). Across all sessions, encoding trial duration correlated significantly with performance ($r_s = -0.42$, $p = 1.35{\cdot}10^{-23}$). **B** As in (**A**), but for the first retrieval trial of a session (n = 499; $F_{4,495} = 79.6$, $p = 1.01{\cdot}10^{-30}$; performance: $t = -12.6$, $p = 8.78{\cdot}10^{-32}$; $ET_i$: $t = 0.28$, $p = 0.778$; across sessions: $r_s = -0.58$, $p = 1.15{\cdot}10^{-46}$). **C** Approach to quantify neural activity ($p_{active}$) within a trial. Step 1: quantify difference score (DS) from 5-second subsample of inferred spiking activity. Step 2: permutation of spike trace, re-quantify DS. The neuron was qualified as active in this subsample when observed DS > 95$^{th}$ percentile of permuted DSs. Step 3: repeat steps 1 and 2 100 times with different baseline and trial subsamples. The sum of active subsamples, divided by 100, yielded the $p_{active}$ for this neuron in this trial. **D** Euclidian distance of the ensemble response vectors, between consecutive encoding trials (ET1, ET2, and ET3) and between ET3 and RT1, sorted by $ET_i$ (consecutive ETs: $F_{2,495} = 3.68$, $p = 0.026$; $ET_i$: $t = -2.70$, $p = 7.85{\cdot}10^{-3}$; mean performance on encoding trials: $t = -0.73$, $p = 0.47$; ET3–RT1: $F_{1,495} = 3.61$, $p = 0.029$; $ET_i$: $t = -2.59$, $p = 0.010$; retrieval trial performance: $t = -0.79$, $p = 0.43$). Filled dots indicate data from each mouse on each session, boxes indicate interquartile range, lines and shaded area indicate mean and SEM, respectively. n.s., non-significant, *** $p < 0.001$.

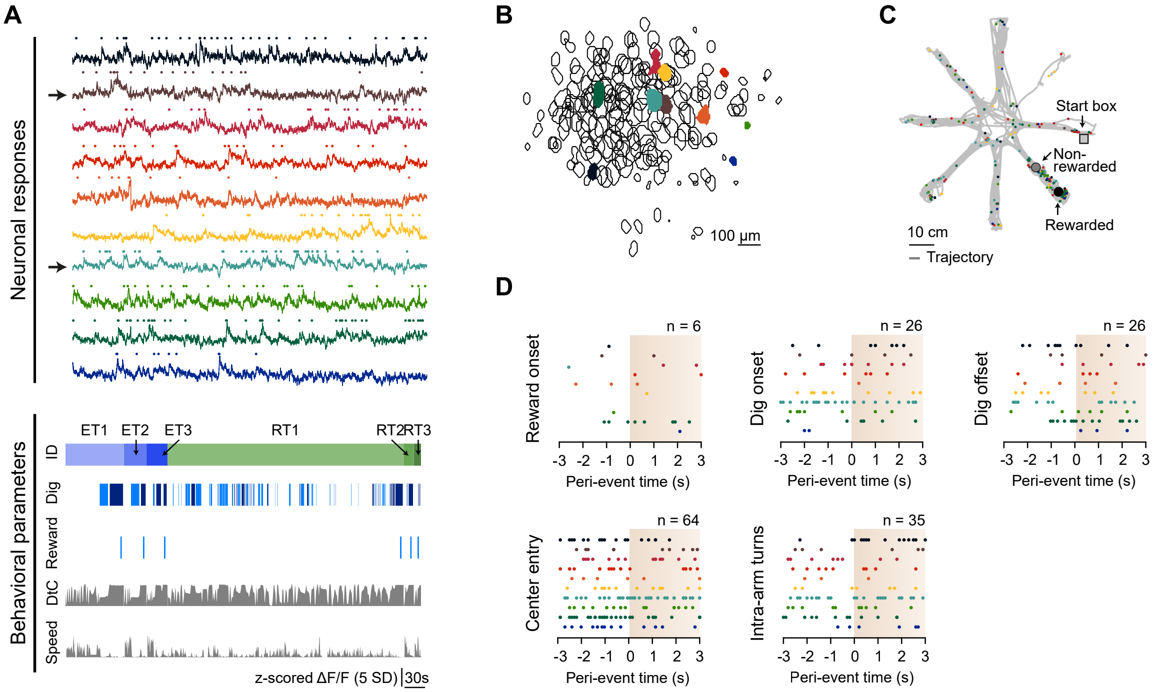


**Supplemental Figure 9.** ***In vivo* calcium imaging of dmPFC neuronal populations in the everyday memory task.** **A** Responses of 10 example neurons (top) and behavioral parameters (bottom) during an example session. Two partially overlapping sources are indicated with an arrow. Behavioral parameters include digging at the rewarded (dark blue) and non-rewarded sandwell (blue), reward consumption ("Reward"), distance to center ("DtC"; max-normalized distance to the central platform), and running speed of the mouse ("Speed"). **B** Outlines of all neurons (black lines) in the field of view of the miniaturized microscope. Example neurons are indicated by color. **C** Locations where the example neurons fired along the mouse's trajectory from the start box via various arms to the non-rewarded sandwell and rewarded sandwell. **D** Peri-event responses of the example neurons, aligned to reward onset (n = 6 events), digging onset (n = 26 events), digging offset (n = 6 events), entry into the center platform (n = 64 events), and intra-arm turns (n = 35 events). Filled dots indicate spikes, color-coded by neuron identity. ID: trial identity SD: standard deviation. Scale bars: 30 s (**A**), 100 µm (**B**), 10 cm (**C**).

**Supplemental Table 1. Overview of habituation procedures across seven days.** Mice freely explored the maze for five minutes with the other cage mates (group) or individually. Mice carried either the dummy or the regular miniaturized microscope, either in the home cage (HC) or in the maze. Mice retrieved a buried chocolate pellet from subsequently larger depths (ranging from 0 to 2 cm below the sand surface) in either in the home cage or in the maze. Pellet retrieval was repeated one, two, or three times.

| | Day 1 | Day 2 | Day 3 | Day 4 | Day 5 | Day 6 | Day 7 |
|---|---|---|---|---|---|---|---|
| Maze | group | group | individual | individual | individual | individual | individual |
| Miniaturized microscope | | dummy (HC) | dummy (maze) | regular (maze) | regular (maze) | regular (maze) | regular (maze) |
| Digging | 2× 0 cm (HC) | 2× 0.5 cm (HC) | 2× 0.5 cm (HC) | 2× 1 cm (HC) | 2× 2 cm (HC) | 1× 2 cm (maze) | 3× 2 cm (maze) |

### Supplemental Methods

#### 1 Mice.

All procedures were performed in accordance with the institutional guidelines of the Max Planck Society and the local government (Regierung von Oberbayern, Germany). Female C57BL/6NRj mice (between 3 and 4.5 months old at the day of surgery, weighing 21.7 ± 1.1 g [mean ± SD]) were communally housed (2–3 mice per cage) in standard, individually ventilated cages, enriched with a running wheel, tunnel and shelter (#13150, #13102, and #13169, Plexx). Mice were kept on an inverted 12-hrs light, 12-hrs dark cycle with lights on at 10 or 11 PM (winter and summer time, respectively) with constant ambient temperature (∼22°C) and humidity (∼55%). Water was always available *ad libitum.* Prior to behavioral experiments, standard chow (#1310, Altromin Spezialfutter GmbH) was available *ad libitum.* From the start of behavioral experiments, mice were food-restricted to 85% of their free-feeding weight [1].

#### 2 Surgical procedures.

**Microprism implant assembly.** The microprism implant was comprised of an aluminum-coated, right-angle microprism (1.5 mm side length, BK7 glass, MPCH-1.5, IMM photonics) that was glued to the center of a circular glass window (3.0 mm diameter coverslip, #1 thickness, BK7 glass, #4100110; Glaswarenfabrik Karl Hecht GmbH; [2]). These components were glued using UV-curing optical adhesive (Norland optical adhesive 71, Norland Products), cured at 50 °C for 12 hours and kept in 70% ethanol until implantation.

**Viral vector injection and microprism implantation.** Mice were anesthetized with a mixture of fentanyl, midazolam and medetomidine in saline (FMM; 0.05 mg $kg^{-1}$, 5 mg $kg^{-1}$, and 0.5 mg $kg^{-1}$; HEXAL AG, Ratiopharm, and Vetoquinol GmbH, respectively, injected intraperitoneally [i.p.]). Sufficient depth of anesthesia was confirmed by absence of the pedal reflex. Mice were placed in a stereotaxic apparatus (Neurostar) equipped with a thermal blanket (Harvard Apparatus). Eyes were covered with a thin layer of ophthalmic ointment (Alcon Pharma GmbH). Lidocaine (10% w $w^{-1}$, AstraZeneca GmbH) was applied onto the scalp for topical anesthesia and carprofen (5 mg $kg^{-1}$, Zoetis, dissolved in saline, injected subcutaneously [s.c]) was administered for analgesia. The skull was exposed, dried, and scraped to facilitate attachment of the head plate. The custom-designed aluminum head plate was fixed in position over the parietal bone using cyanoacrylate glue (Ultra Gel Matic, Pattex) and subsequently secured with dental acrylic (Paladur, Kulzer GmbH) mixed with black pigment (47150, Kremer Pigmente).

A 3 mm circular craniotomy was created centered at anteroposterior (AP) 2.0 mm and mediolateral (ML) 0.75 mm from bregma over the implantation hemisphere. The brain was kept covered in cortex buffer (125 mM NaCl, 5 mM KCl, 10 mM glucose, 10 mM HEPES, 2 mM $CaCl_2{\cdot}2H_20$, and 2 mM $MgSO_4{\cdot}7H_2O$) throughout.

To label a sparse population of dmPFC excitatory neurons, mice were injected with a mixture of two adeno-associated viral vectors (AAVs): AAV2/1 CaMKII0.4-Cre ($4.6{\cdot}10^9$ GC $ml^{-1}$, gift from James M. Wilson, Addgene plasmid # 105558, LOT CS0843, Penn Vector Core) and AAV2/1 hSyn-flex-GCaMP6m ($3.2{\cdot}10^{12}$ GC $ml^{-1}$, gift from Douglas Kim & GENIE Project, Addgene plasmid # 100838, Penn Vector Core). We targeted three injection sites (ML 0.3 mm, dorsoventral [DV] -1.6 mm) along the anteroposterior axis (AP 1.6 mm, 2.0 mm, and 2.4 mm). At each injection site, 150 nl of the viral vector mixture was injected using a pulled and beveled glass pipette (30 $\mu$m outer diameter at the tip, 1408472, Hilgenberg GmbH) at an injection speed of 25 nl $min^{-1}$. The glass pipette was slowly retracted 10 min after the end of the injection.

To implant the microprism, a durotomy was performed over the implantation hemisphere using a 30 G needle and fine-tipped forceps [2]. The dura remained intact over the contralateral (imaged) hemisphere. The microprism was manually lowered into the sagittal fissure, with the front face of the microprism flush with the contralateral hemisphere. The glass window of the implant was placed ∼100 µm below the inner surface of the skull, lightly compressing the dorsal cortical surface. The microprism was bonded to the skull using a thin layer of cyanoacrylate glue and subsequently secured with black pigmented dental acrylic. After surgery, anesthetic agents were antagonized with a mixture of naloxone, flumazenil, and atipamezole in saline (NFA; 1.2 mg $kg^{-1}$, 0.5 mg $kg^{-1}$, and 2.5 mg $kg^{-1}$, Ratiopharm, HEXAL AG, and Prodivet pharmaceuticals, respectively, injected s.c.). Mice were placed under a heat lamp to recover and were then returned to their home cage. Mice received carprofen (5 mg $kg^{-1}$, injected s.c.) and dexamethasone (2 µg $kg^{-1}$, Sigma, injected s.c.) for the two subsequent days. Implantation of the microprism resulted in mild microgliosis three months post-op without introducing long-term astrogliosis or apoptosis, comparable to what was observed after a cranial window implant (**Supplemental Figure 7A–E**) [3].

**Miniaturized microscope lens placement.** Two weeks after microprism placement, mice were anesthetized with FMM, and sufficient depth of anesthesia was confirmed by absence of the pedal reflex. The miniaturized microscope, mounted on a rigid holder (Doric Lenses) and attached to a XYZ translation stage (Luigs Neumann), was lowered over the microprism implant until neurons came sharply into view. The center of the field of view was approximately at AP 2.2 mm (± 250 µm), DV -1.0 mm (± 250 µm) from bregma, allowing imaging of upper layer 2/3. The miniaturized microscope lens and the affixed adjustment ring were bonded to the overlying dental cement using a thin layer of cyanoacrylate glue and subsequently secured with dental acrylic mixed with black pigment. The miniaturized microscope body was removed from the lens, mice were injected with NFA and placed under a heat lamp to recover.

#### 3 Behavioral procedures in the everyday memory task.

**Apparatus**. Behavioral training on the everyday memory task [4] was performed in a custom-made, 8-arm radial maze (**Figure 1A, S1A**). Arms (30 × 10 cm) radiated from the center of the maze (18 cm diameter) and were termed “cardinal” when parallel to the room’s walls (**Figure S1A**). Arms were flanked with clear Plexiglas walls (25 cm height) and contained cutouts for reward cups. The cutouts measured 5 cm in diameter and were centered 5.5 cm and 16.5 cm from the end of the arms. During behavioral experiments, cutouts contained either a sandwell (4 cm inner diameter, 4 cm depth; **Figure S1B**) or were covered by a white Plexiglas lid. The sandwells could not be seen from a distance from the mouse’s perspective. The sandwells were subdivided in a center and surround compartment using semi-circular, perforated 3D-printed removable mesh cups (**Figure S1B, C**). The surround compartment contained 20 chocolate-flavored pellets (190 mg, chocolate flavor, Dustless Precision Pellet, Bio-Serv; **Figure S1B**) that served as masking odors and that were inaccessible to the mouse. The rewarded sandwell contained one additional accessible chocolate-flavored pellet placed in the center compartment, 2 cm below the sand surface (**Figure S1B, C**). A start box (10 × 11 × 25 cm, L × W × H) with black Plexiglas walls was mounted at the end of one of the cardinal arms. The start box was equipped with a pneumatic door, which could be opened and closed remotely. The room contained 3D cues (multi-colored paper decorations in various shapes and sizes; **Figure S1A**) and two proximal landmarks (LEGO® DUPLO® tower and plush dice; **Figure S1A**). The illumination of the laboratory room was maintained at a moderate level by LED lamps emitting white light. Mouse position was recorded using an overhead webcam (LifeCam Studio, Microsoft) and tracked with custom-written Python routines.

**Habituation and pre-training.** All behavioral training was conducted during the active (night) cycle of the mouse, approximately between 12:00 and 19:00 o’clock. After mice were transported to the behavioral training room, they were weighed and kept in their home cage for a minimum of 20 min prior to experimentation.

During the first 7 days of the experiment, the mice were habituated to the maze, habituated to carrying the miniaturized microscope, and trained to dig for a chocolate pellet in the sandwells (**Table S1**).

To habituate the mice to the maze, all cage mates were placed together in the maze on habituation day 1 and 2 and explored the maze for 5 min. Mice were habituated individually on day 3–7 for 5 min each. On habituation day 1–5, mice were initially placed onto the central platform. On habituation day 6–7, mice were initially placed into the start box, which was opened after 30 s, after which mice were able to retrieve a buried chocolate pellet. After finding the pellet, mice were gently guided back to the start box, where they consumed the pellet. On habituation day 7, two non-rewarded sandwells were also present in the maze.

To habituate the mice to carry the miniaturized microscope, they were briefly manually head-fixed and either the dummy miniaturized microscope (habituation day 2–3) or the actual miniaturized microscope (habituation day 4–7) was affixed to the imaging cannula. Subsequently, mice either freely explored the home cage (habituation day 2) or the maze (habituation day 3–5) for 5 min. On the final two days of habituation, the mice carried the miniaturized microscope while digging for the reward in the maze.

To train the mice to dig in the sandwells for a food reward, a chocolate pellet was placed progressively deeper into a sandwell placed in the home cage (habituation day 1–5). Mice had to retrieve the pellet and consume it completely before a second pellet was buried at the same depth.

**Main behavioral training phase.** Each session consisted of three encoding trials (ETs) and three retrieval trials (RTs; **Figure 1B, Video 1**). The encoding trials had an intertrial interval (termed the "encoding intertrial interval", abbreviated $ET_i$) of either 30 s ("massed"), 10 min, 30 min or 60 min (all "spaced"). The delay between the final encoding trial and the subsequent retrieval or probe trial was 2.5 hrs or 24 hrs (termed the "retrieval delay", abbreviated $RT_d$, **Figure 1B**). All eight combinations of $ET_i$ and $RT_d$ formed one session block and blocks were repeated either three (n = 10 mice) or five (n = 10 mice) times (**Figure 1D**).

At the start of an encoding trial, the mouse was placed into the start box for 60 s. Subsequently, the experimenter would remotely open the start box and the mouse could explore the maze containing two sandwells, the rewarded sandwell and the non-rewarded sandwell. The location of both these sandwells was kept constant throughout a single session. Once a mouse found the buried pellet, the mouse was gently nudged and ran back to the start box. The start box was remotely closed and the mouse consumed the reward there. In sessions with an $ET_i$ of 30 s, the door of the start box was opened after 30 s and two more encoding trials were conducted in the same manner. At the end of the final $ET_i$, the mouse was kept in the start box for 60 s and subsequently placed back in its home cage. In sessions with an $ET_i$ longer than 30 s, mice were placed back in the home cage after 60 s and remained there during the $ET_i$. Retrieval trials were conducted either 2.5 or 24 hrs after completion of the third encoding trial. In retrieval trials, the maze contained six sandwells: the rewarded sandwell, the non-rewarded sandwell, and four unfamiliar non-rewarded sandwells ("non-cued sandwells"). Training was carried out the same way as in encoding trials, except that the interval between subsequent retrieval trials was kept constant at 30 s. The next session was started on the following day, creating a non-training period of between 17 and 22 hrs, depending on the $ET_i$ and $RT_d$ of the session.

The first retrieval trial was occasionally replaced by a probe trial (**Video 2, 3**). A probe trial was conducted as a regular retrieval trial with the notable exception that the rewarded sandwell did not contain a chocolate pellet for the first 60 s after the opening of the start box. After 60 s, the experimenter entered the maze room and placed one chocolate pellet in the sandwell, which was retrieved by the mouse as described above. Subsequently, two additional retrieval trials were conducted. Probe trial sessions were interleaved with generally five but minimally two non-probe trial sessions.

**Additional behavioral interventions.** After conclusion of the main behavioral training phase, several control experiments were conducted. These control sessions were generally conducted similar to those in the main experimental phase, but concluded with a probe trial and with the following alterations:

*Single encoding trial.* To evaluate whether mice were able to recall the rewarded sandwell location after a single encoding trial ($RT_d$ 2.5 hrs), two slightly different versions of a "single encoding trial" experiment were conducted (**Figure S2A**). In one version, the mouse was presented with two additional chocolate pellets in the start box immediately after the encoding trial ended and then returned to the home cage for the remainder of the $RT_d$ . In the other version, we controlled for context exposure by re-introducing the mouse twice (interval 30 min) into the start box of the maze, where it received a chocolate pellet.

*Incongruent training.* To evaluate whether mice retrieved memory encoded during each encoding trial equally and confirm the absence of primacy or recency effects, the location of the rewarded sandwell was changed after each encoding trial ($ET_i$ 30 s, $RT_d$ 2.5 hrs; **Figure S2D**). Enhanced learning during a specific encoding trial was hypothesized to result in increased digging at the rewarded sandwell of that encoding trial.

*Egocentric navigation strategy.* To evaluate whether mice used an egocentric navigation strategy, the start box location was altered during the retrieval delay ($ET_i$ 30 s or 30 min, $RT_d$ 24 hrs; **Figure S3A**). If animals followed an egocentric strategy, they were hypothesized to retrace the previously learned path in the probe trial, leading them to a non-cued sandwell, termed the "egocentric sandwell".

*Allocentric navigation strategy.* To evaluate whether mice could remember the rewarded sandwell location when forced to use an allocentric navigation strategy, the start box was located at a different arm in each ET, changing the path from start box to rewarded sandwell ($ET_i$ 30 min, $RT_d$ 24 hrs; **Figure S3C**).

*Extended encoding intertrial interval.* This session was conducted with an encoding intertrial interval of 180 min and a retrieval delay of 24 hrs (**Figure S4**).

*Extended retrieval delay.* This session was conducted with an encoding intertrial interval of 30 s and a retrieval delay of 48 hrs (**Figure S4**).

**Trial randomization.** The entire behavioral experiment was subdivided into blocks of eight sessions, comprising all combinations of $ET_i$ (4 durations) and $RT_d$ (2 durations; **Figure 1D**). Within a block, the session's $ET_i$ and $RT_d$ were randomly distributed. The start box location was changed between sessions, as well as the order in which the mice were tested. For a given mouse, the location of the rewarded sandwell was randomized across sessions, so neither the same sandwell location, nor the same egocentric path to the sandwell, would be rewarded on two consecutive sessions (**Figure S1D**). The experimenter was not blinded to the experimental condition.

**Control for olfactory cues.** Multiple precautions were put in place to minimize olfactory cues. The sand that filled the sandwells (Prestige Muschelsand Kristal, Versele Laga) contained 5% w $w^{-1}$ Garam Masala powder. It was prepared fresh daily and kept in an airtight container. Twenty chocolate pellets were placed below the mesh divider in the sandwell, making them inaccessible to the mouse while allowing transmission of chocolate odors through the perforated partition wall (**Figure S1C**). On each trial, the rewarded sandwell contained only one additional pellet, placed at the bottom (2 cm from surface) of the sandwell. In-between trials, all sandwells were refilled, irrespective of whether the mouse had dug at the sandwell. Any sand on the maze was brushed and vacuumed away, and all arms were carefully wiped with 40% ethanol. At the end of each day, the maze was thoroughly cleaned using 80% ethanol. The maze was rotated by 90° on a weekly basis.

**Quantification.** For each encoding and retrieval trial, the parameters "latency" and "error" were manually recorded. Latency was defined as the time from door opening to the start of the final digging period before retrieving the reward. Error was defined as the number of sandwells the mouse dug at before retrieving the reward. Of note, multiple dig periods at the same incorrect sandwell were scored as one error. For each trial, the performance index (PI) was calculated as

$$PI = \frac{error_{\text{max}} - error_{\text{observed}}}{error_{\text{max}}} \cdot 100\%$$

with $\text{error}_{\text{max}} = 1$ for encoding trials and $\text{error}_{\text{max}} = 5$ for retrieval trials. Retention was quantified as the relative difference in the mean PI of RT1 and ET3 for each individual mouse, across sessions of the same encoding intertrial interval and retrieval delay. For each probe trial, the time in the rewarded, non-rewarded and non-cued arms was automatically recorded using custom-written Python routines. The behavioral videos were annotated frame-by-frame for position, speed, and distance to the nearest sandwell using custom-written MATLAB (version R2016b; MathWorks) routines. Frames with mouse positions $< 1$ cm from a sandwell and movement $< 0.4$ cm $s^{-1}$ were labeled as dig frames. TIn probe trials, the relative time spent digging at the rewarded sandwell as compared to the total dig time at both the rewarded and non-rewarded sandwell yielded the occupancy difference score (ODS):

$$ODS = \frac{dig_{\text{rewarded}}}{dig_{\text{rewarded}} + dig_{\text{non-rewarded}}}$$

An occupancy difference score of 0 indicated equal dig time at the rewarded and non-rewarded sandwell, positive and negative values indicate more digging at the rewarded and non-rewarded sandwell, respectively.

#### 4 Chemogenetic silencing.

**Surgical Procedures.** Cre-dependent AAVs were used to express either the control fluorophore mCherry (AAV 2/9 hSyn-DIO-mCherry, $2.1 \cdot 10^{12}$ GC ml$^{-1}$, gift from Bryan Roth, Addgene viral prep # 50459-AAV9, LOT v49659) or the Gi-coupled receptor hM4D(Gi) conjugated to the fluorophore mCherry (AAV 2/9 hSyn-DIO-hM4D(Gi)-mCherry, $2.3 \cdot 10^{12}$ GC ml$^{-1}$, gift from Bryan Roth, Addgene viral prep # 44362-AAV9, LOT v54501). A viral vector mixture of an AAV encoding a Cre recombinase (AAV2/1 CaMKII0.4-Cre, $2.1 \cdot 10^{11}$ GC ml$^{-1}$, gift from James M. Wilson, Addgene viral prep # 105558-AAV1, LOT CS0843) and either the control fluorophore or hM4D(Gi)-encoding AAV was bilaterally injected at two locations along the anteroposterior axis in the dmPFC (+2.5 mm AP, ±0.3 mm ML, -1.0 mm DV, and +1.5 mm AP, ±0.3 mm ML, -2.0mm DV, relative to bregma) using a pulled and beveled glass pipette (30 $\mu$m outer diameter). Each injection contained 150 nl of the viral vector mixture, was injected at 30 nl min$^{-1}$, and was flanked by a 5 min pre- and post-injection window. Behavioral and slice electrophysiological experiments were started after a minimum of 2 and 4 weeks after surgical procedures, respectively.

***Ex vivo* electrophysiology.** Mice were decapitated under isoflurane anesthesia, the brains were dissected out within 3 min and the hemispheres were separated. Coronal slices (300 µm) containing the dmPFC were prepared using a vibratome (VT1200S, Leica) while the brain remained submerged in ice-cold "cutting" aCSF (85 mM NaCl, 75 mM sucrose, 2.5 mM KCl, 25 mM glucose, 1.25 mM $NaH_2PO_4$, 4 mM $MgCl_2$, 0.5 mM $CaCl_2$, and 24 mM $NaHCO_3$; 320–325 mOsm; carbogenated 95% $O_2$, 5% $CO_2$ vol vol$^{-1}$). Slices were transferred to a custom, light-shielded slice incubation chamber containing carbogenated cutting aCSF and allowed to equilibrate for 30 min at 37 °C. After the initial incubation, slices were transferred to a custom, light-shielded carbogenated slice incubation chamber containing "recording" aCSF (127 mM NaCl, 2.5 mM KCl, 10 mM glucose, 1.25 mM $NaH_2PO_4$, 2 mM $MgCl_2$, 2 mM $CaCl_2$, and 26 mM $NaHCO_3$; carbogenated 95% $O_2$, 5% $CO_2$ vol vol$^{-1}$) at room temperature for up to 12 hrs.

For recordings, a slice was transferred to a recording chamber under continuous perfusion of carbogenated recording aCSF (1 ml min$^{-1}$). The dmPFC was identified under visual guidance of a fluorescence microscope (BX51WI, Olympus). Neurons expressing mCherry were identified by illuminating the slice with red light using a mercury-vapor lamp and a filter (excitation 560/40 nm, emission 630/75 nm, 49008, Chroma). We performed intracellular patch-clamp recordings using electrodes (3–5 MΩ) filled with K-gluconate-based internal solution (135 mM K-gluconate, 0.2 mM EGTA, 10 mM HEPES, 4 mM $MgCl_2$, 4 mM $Na_2$-ATP, 0.4 mM Na-GTP, 10 mM $Na_2$-phosphocreatine, 3 mM ascorbate, and 50 nM Alexa 488, pH 7.2, 295–300 mOsm). Electrical signals were acquired using an amplifier (AxoClamp-2B, Axon Instruments, Inc.), post-amplified (440, Brownlee), low-pass filtered (3 kHz cut-off), noise-filtered (HumBug Noise Eliminator), digitized at 10 kHz, and recorded using custom-written LabVIEW routines (National Instruments).

To determine the effect of clozapine-$N$-oxide (CNO, HB6149, HelloBio) on intrinsic excitability and current-evoked excitability of dmPFC excitatory neurons, two current injection protocols were executed. First, to establish the relationship between the injected current and evoked potential, a step protocol was executed (step: -100 pA prepulse injection for 100 ms, followed by a 500-ms delay, followed by a current injection ranging from -450 pA to 450 pA at steps of 50 pA for 750 ms; each step separated by 10 s). Second, to determine the rheobase of the neuron, a step protocol with higher resolution was executed (step: -100 pA prepulse injection for 100 ms, followed by a 500-ms delay, followed by a current injection ranging from 0 pA to 150 pA at steps of 10 pA for 750 ms; each step separated by 10 s). To assess the influence of CNO on the resting membrane potential, the resting membrane potential was measured at 1 Hz intervals starting 5 min prior to the influx of recording aCSF containing 50 µM CNO until 10 min post-influx. Subsequently, both aforementioned step protocols were performed again. Analysis of the recorded traces was performed using custom-written LabVIEW and MATLAB routines. Only neurons whose series resistance did not change more than 20% during the recording were included in the final analysis.

**Behavioral chemogenetic experiments.** Behavioral experiments followed a full factorial $2^4$ design, with the four factors protein expression (mCherry or hM4D(Gi)-mCherry conjugate), injection substance (saline [vehicle] or CNO in saline), injection time point (before an encoding trial or before a retrieval trial), and the session's $ET_i$ (0.5 min or 60 min). Behavioral training was executed as described in section 2. Injections (i.p.) with either vehicle or CNO (5 mg $kg^{-1}$) were performed 45 min before behavioral testing. Injections were administered between 38 and 71 days after viral vector injection. Sessions with injections were interleaved with at least two sessions without injections.

#### 5 Miniaturized microscopy.

**Image acquisition.** The training period during which in vivo calcium imaging was performed lasted up to 2.5 months for each mouse. Images were acquired with a commercially available miniaturized microscope (Basic Fluorescence Microscopy System - Surface, Doric Lenses) at a frame rate of 10 Hz and a resolution of 630 × 630 pixels (field of view 1 $mm^2$). Laser power under the objective lens (2× magnification, 0.5 NA) was <1 mW for all imaging experiments. The excitation wavelength was 458 nm. Imaging typically lasted 3 min per behavioral trial, and never lasted more than a total of 20 min per imaging session. Before the start of imaging, the mouse was briefly held head-fixed by the experimenter to attach the miniaturized microscope. The imaging location was verified based on landmarks such as blood vessels, and typically did not drift strongly over time. The miniaturized microscope was always removed before the mouse was returned to the home cage.

**Image registration and source extraction.** iming of individual behavioral video and miniaturized microscopy frames was synchronized using data acquisition cards (PCLe 6321, National Instruments). Behavioral data were downsampled to fit the miniaturized microscope frame acquisition rate. Imaging frames were spatially downsampled to 256 × 256 pixels. Frames within a single recording were registered to each other to correct for motion artefacts using the NoRMCorre package [5]. Subsequently, all registered frames that were collected over the course of a single session were concatenated into a single stack and aligned again using the NoRMCorre package [5]. Single neuron $Ca^{2+}$ activity traces were extracted from the fluorescent imaging time series by applying constrained nonnegative matrix factorization for microendoscopic data (CNMF-E; [6]). Putative sources that did not adhere to two inclusion criteria were removed from the dataset. First, temporal transients of individual sources were fit with a single-term exponential and any sources whose fitted exponential decay factor was below -0.07 were removed. Second, neurons that had less than six transients during a session were removed. On average, we included 210 ± 99 neurons (mean ± SD) per session. All subsequent analyses were conducted using the deconvolved spike rate generated by the CNMF-E algorithm.

**Quantifying behavioral modulation of neuronal responses using a generalized linear model.** A generalized linear model (GLM) was fitted to the spiking activity of single neurons to establish the influence of specific behavioral parameters (task predictors; *tp*s) on neuronal activity (similar to approaches taken in [7]; **Figure 4A**). The model incorporated task predictors relating to reward, motor activity, and decision-making. Reward-related *tp*s included reward onset and reward anticipation (i.e. final entry into the rewarded arm), motor activity-related *tp*s included onset and offset of digging at a sandwell, running speed, and running acceleration, and decision-related *tp*s included entry into the central platform and intra-arm reversals of heading direction.

The task predictors were processed as follows. Continuous task predictors, i.e. running speed and running acceleration, were binned into 500 ms bins. Categorical task predictors, i.e. reward onset, reward anticipation, onset and offset of digging at a sandwell, entry into the central platform, and intra-arm reversals, were represented as boxcar functions that were set to the value one at the time of onset and zero everywhere else. Next, categorical predictors were convolved with five evenly spaced Gaussian basis functions (1.4 s half-width at half-height). The third Gaussian was centered on the predictor onset and the peaks were spaced 2.5 s apart. Versions of the GLM with different numbers of Gaussian basis functions and different parameter values were also fitted, and yielded qualitatively similar results. Finally, all predictors were rescaled to the range [0,1]. These manipulations resulted in a design matrix comprised of a constant predictor and 32 task predictors.

The neuron's inferred binarized spiking activity was downsampled to 2 Hz and split into a training dataset (70% of frames, randomly selected) and a test dataset (the remaining 30% of frames). The training set was used to fit a Bernoulli GLM. To exclude non-informative task predictors from the final GLM, the MATLAB function "*lasso*" regularized the task predictors with the additional specifiers "*NumLambda*" and "*CV*" set to "10" and "10", respectively. Only regressors with non-zero regression coefficients at the minimum-deviance point were retained in the final, regularized model. The obtained regression coefficients were multiplied with the task predictor values and summed across temporally offset predictors of the same underlying task predictor. This resulted in a design matrix comprised of the constant predictor and eight deconvolved task predictors. The absence of multicollinearity was verified by calculating the variance inflation factor, using a cut-off of 4. The regularized and deconvolved design matrix and the training dataset were supplied to the MATLAB function "*fitglm*", with the additional specifiers "*modelspec*", "*Distribution*", and "*link*" set to "linear", "binomial", and "logit", respectively. When a resulting regression coefficient was significant on the t-test after Bonferroni correction, the neuron was labeled "modulated" by this task predictor.

To quantify model fit, the GLM was fitted to a permuted spike trace using the previously defined task predictors. Model fit was quantified as the adjusted $R^2$ of the model fitted to the observed spiking trace. The model's decoding performance was quantified by comparing the observed test dataset with a decoded test data set, which was obtained using the MATLAB function "*predict*".

**Quantification of neuronal activity spanning complete trials.** To quantify the activity of a neuron during a trial, we used a probabilistic measure ($p_{active}$; **Figure S8C**). Trials were subdivided into the baseline and trial period. The baseline period was the 60-second period that the mouse spent in the start box prior to maze exploration. The trial period was the period from the first entry into the maze until 2 s after the mouse had retrieved the reward. Only trials with a minimum duration of 10 s were included in these analyses. To calculate $p_{active}$, we randomly selected a continuous, 5-second subsection from the baseline and trial period, thereby controlling for trial duration. The instantaneous inferred spike rate during both the baseline and trial subsection was averaged across the subsection. The average baseline inferred spike rate was subtracted from the average trial inferred spike rate, yielding the observed "trial activity rate". Subsequently, the trial activity rate was calculated 1000× using permuted spike rate data, and the resulting permuted trial activity rates were stored in a 1000 × 1 vector. If the observed trial activity rate was larger than the $95^{th}$ percentile of this vector, the neuron was labeled "active" for this particular 5-second subsection. The procedure outlined above was repeated 100 times for different, pseudo-randomly selected (i.e. non-duplicate) sections of baseline and trial periods. $p_{active}$ was defined as the fraction of these 100 subsamples in which the neuron was labeled "active". This procedure was repeated for all neurons and all trials, and the $p_{active}$ values were concatenated into an N × 1 vector (N = neurons), which was termed the ensemble response vector. We calculated the Pearson correlation coefficient between the ensemble response vectors of the six trials in a session, which yielded the ensemble correlation matrix.

#### 6 Histology and immunohistochemistry.

**Perfusion.** Mice were deeply anesthetized using FMM and transcardially perfused with saline containing lidocaine and heparin (25 mg ml$^{-1}$ and 14 mg ml$^{-1}$, respectively) for 5 min at a flow rate of 3 ml min$^{-1}$, followed by 4% PFA in PBS for 5 min at a flow rate of 3 ml min$^{-1}$. Brains were dissected out and post-fixed in 4% PFA in PBS for at least one week at 4 °C, followed by cryoprotection with 30% sucrose in PBS for 3 days. Coronal sections (40 $\mu$m) were cut on a sliding microtome (HM 400, Thermo Fisher Scientific) and were kept free-floating in PBS at 4 °C until further processing.

**Immediate early gene expression experiments.** Mice were pseudo-randomly assigned to one of six groups, ensuring that cohoused mice were equally distributed across conditions. These groups underwent behavioral training using an $ET_i$ of 30 s, 10 min, 30 min, or 60 min, were placed in the maze for 1 min without training ("handled control") or were merely handled ("home cage control"). Mice were perfused 90 min after the start of ET2 ($ET_i$ 30 s, 10 min, and 30 min), 90 min after the middle of the interval between ET2 and ET3 ($ET_i$ 60 min), or 90 min after handling (handled and home cage control). After perfusion, immunohistochemistry was carried out using the primary antibody rabbit anti-c-Fos (1:1000; 226 003, Synaptic Systems). After washing, sections were incubated with a species-specific secondary antibody conjugated to Cy3 (1:200; 111-165-045, dianova) and mounted with mounting medium containing DAPI (H-1200, Vectashield). Five serial optical sections of the dmPFC (spaced at 1 $\mu$m) were acquired using a laser-scanning confocal microscope (TCS SP8, Leica) with a 20× objective (NA 0.75). Images had a resolution of 1024 × 1024 pixels and color channels were acquired sequentially using excitation lasers for DAPI (excitation at 405 nm, emission at 410–419 nm) and Cy3 (excitation at 561 nm, emission at 575–714 nm).

**Validation of microprism placement.** After perfusion, sections were mounted with mounting medium containing DAPI and the location of the microprism implant and viral vector transduction into the dmPFC were verified using a fluorescence microscope (Axio Imager 2, Carl Zeiss Microscopy) and compared to a reference atlas [8].

**Analysis of injury and inflammation.** Mice that were implanted with a microprism were perfused 3 months post-operation. Immunohistochemistry was carried out using the primary antibodies rabbit anti-Iba1 (1:1000; 019 19741, Wako) and chicken anti-GFAP (1:600; ab4674, Abcam). After washing in PBS, sections were incubated with a species-specific secondary antibody conjugated to Cy3 (1:200; 111-165-045, dianova) or Alexa 647 (1:200; A21449, Thermo Fisher) and mounted with mounting medium containing DAPI. A subset of sections was stained using Red TUNEL stain following the manufacturer's instructions (Click-iT™ Plus TUNEL Assay, Thermo Fisher Scientific). Appropriate positive and negative controls were carried out for all stains. Images were acquired using a laser-scanning confocal microscope.

**Image quantification.** Quantitative analyses were performed within a region of interest (0.31 $mm^2$ per hemisphere) in the dmPFC (∼4 slices per brain) using the "Cell COunter" plug-in for ImageJ [9] by counting the number of TUNEL-labeled, c-Fos immunopositive, GFAP immunopositive, or Iba1 immunopositive neurons.
